# TORC1-driven translation of *Nucleoporin44A* promotes chromatin remodeling and germ cell-to-maternal transition in Drosophila

**DOI:** 10.1101/2025.03.14.643309

**Authors:** Noor M. Kotb, Gulay Ulukaya, Anupriya Ramamoorthy, Lina Seojin Park, Julia Tang, Dan Hasson, Prashanth Rangan

## Abstract

Oocyte specification is a critical developmental transition that requires the coordinated repression of germ cell-specific genes and activation of the maternal program to support embryogenesis. In Drosophila, the timely repression of germ cell and early oogenesis genes is essential for this transition, yet the mechanisms that coordinate this process remain unclear. Here, we uncover an unexpected translation-chromatin axis, where transient Target of Rapamycin Complex 1 (TORC1)-driven translation triggers chromatin remodeling, ensuring irreversible oocyte fate commitment. Through a screen, we identified ribosome biogenesis regulators, including Zinc finger protein RP-8 (Zfrp8) and TORC1 components, as key mediators of gene silencing. We show that TORC1 activity increases during oocyte specification, and disrupting ribosome biogenesis, translation, or TORC1 function prevents proper heterochromatin formation, leading to epigenetic instability. Polysome-seq analysis of *zfrp8*-depleted ovaries revealed that Zfrp8 is required for the translation of *Nucleoporin 44A* (*Nup44A*), a key nuclear pore complex (NPC) component. Given the role of the NPC in chromatin organization, independent disruption of *Nup44A* results in defective silencing of the germ cell and early oogenesis genes. Our findings reveal a mechanism in which translation-driven NPC remodeling coordinates heterochromatin establishment, facilitating the germ cell-to-maternal transition and ensuring proper oocyte fate commitment.

## Introduction

The formation of gametes is essential for sexual reproduction. During oogenesis, germline stem cells (GSCs) or germ cells differentiate and undergo meiosis to produce mature oocytes (Spradling et al., 2011; Lehmann, 2012). Once specified, the oocyte accumulates maternal RNAs, which are crucial for supporting early embryonic development (Tadros and Lipshitz, 2009; Laver et al., 2015). The mechanisms underlying the transition from a GSC or germ cell-specific transcriptional program to a maternal transcriptional program remain poorly understood.

Drosophila oogenesis is marked by a well-characterized transition from GSCs to an oocyte. The GSCs reside in the germarium, where they undergo asymmetric division to produce one GSC and a cystoblast (CB) (**Figure 1A-A1**) (Spradling et al., 2001; Xie and Spradling, 1998). CBs differentiate by expressing *bag of marbles* (*bam*) and undergo four incomplete mitotic divisions, forming 2-, 4-, 8-, and eventually 16-cell cysts (McKearin and Ohlstein, 1995; Ohlstein and McKearin, 1997). GSCs and CBs are marked by spectrosomes, which transition into elongated fusomes in developing cysts (**Figure 1A-A1**) (Lin et al., 1994; de Cuevas et al., 1996; de Cuevas and Spradling, 1998). At the 16-cell cyst stage, one cell becomes the oocyte, while the remaining 15 cells develop into nurse cells that support the growing oocyte and produce maternal components (Huynh and St Johnston, 2004). The 16-cell cyst is then encapsulated by somatic follicle cells to form an egg chamber that develops into a mature egg. (**Figure 1A-A1**) (Nystul and Spradling, 2010).

**Figure 1.**
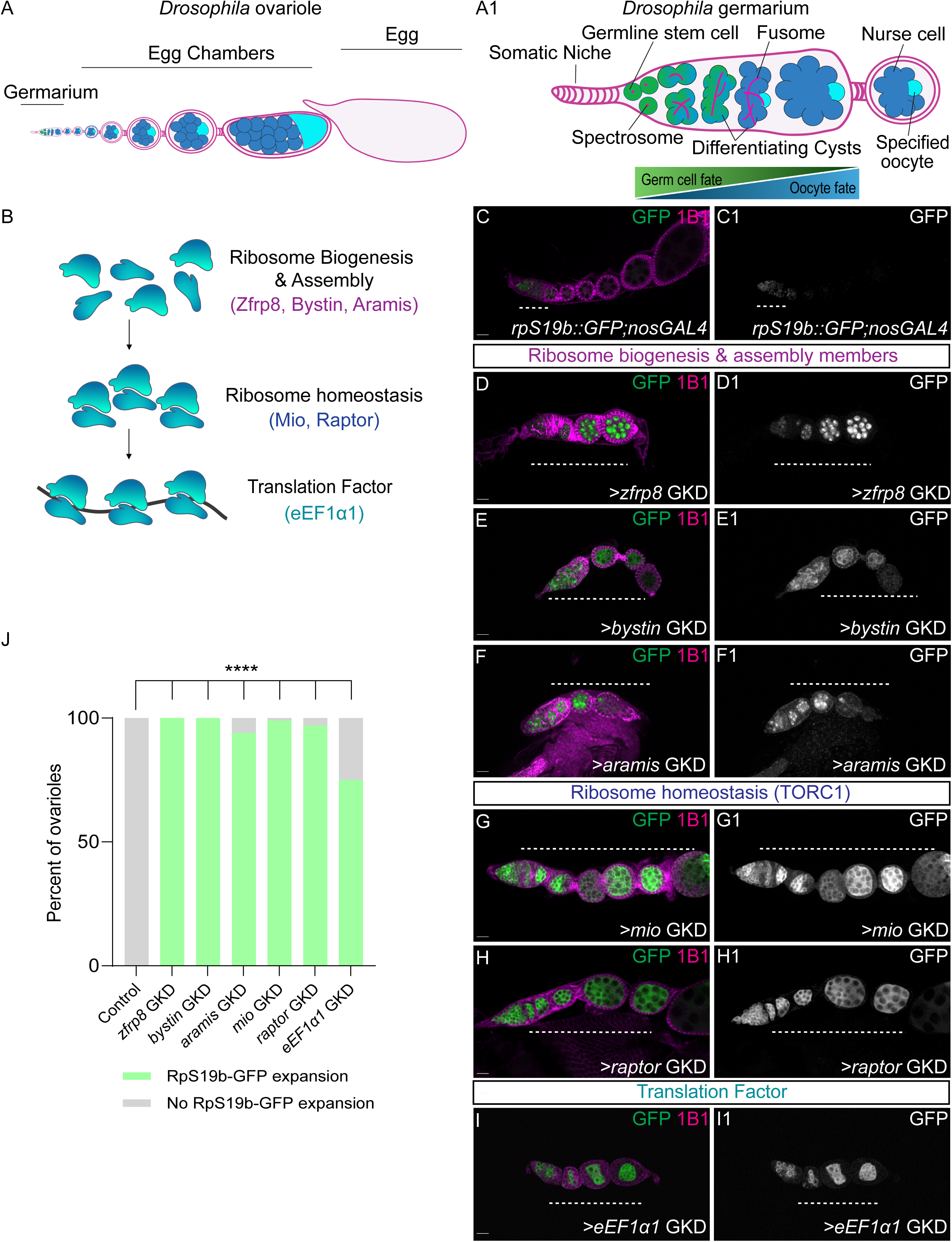
Ribosome level regulators are required for silencing the rpS19b reporter during oogenesis. (A) A schematic of a *Drosophila* ovariole consisting of a germarium containing early stages (green), egg chambers (blue) surrounded by somatic cells and an egg. Egg chambers grow and produce an egg (light blue). (A1) A schematic depicting the *Drosophila* germarium. Near the somatic niche (light pink), are the germline stem cells (GSCs; green), dividing to give rise to daughter cells called cystoblasts (CBs). Both GSCs and CBs are identified by spectrosomes (red). CBs differentiate, giving rise to 2-, 4-, 8-, and 16-cell cysts (blue), marked by fusomes (red). Within the 16-cell cyst, a single cell initiates meiosis and becomes the oocyte (dark blue), while the remaining 15 cells develop into nurse cells (light blue). Genes associated with early oogenesis are active in the undifferentiated stages but decrease in expression as the oocyte is specified. Conversely, maternal genes become enriched in the later, differentiated stages. (B) Schematic of the genetic screen targeting regulators of ribosome biogenesis (zfrp8, bystin, and aramis), ribosome homeostasis (mio and raptor), and translation (eEF1α1) to assess their role in gene silencing. (C–C1) Ovariole of control *rpS19b::GFP;nosGAL4* (C) and grayscale (C1) stained for GFP (green) and 1B1 (magenta). In control, GFP is expressed in the undifferentiated stages and early cysts (white dashed line) and then silenced. (D-I1) Ovarioles from ribosome regulator germline knockdown (GKD) lines stained for GFP (green) and 1B1 (magenta). Grayscale panels (D1-I1) show GFP expression. Loss of *zfrp8* (D-D1), *bystin* (E-E1), *aramis* (F-F1), *mio* (G-G1), *raptor* (H-H1), and *eEF1α1* (I-I1) resulted in ectopic expression of RpS19b::GFP (white dashed line), failure of egg chamber growth, and oogenesis defects. (J) Quantification of ovarioles with expanded RpS19b::GFP expression. Statistics: Two-tailed t-test; n = 50 ovarioles per genotype. ns, p > 0.05; *p < 0.05; **p < 0.01; ***p < 0.001; ****p < 0.0001. Scale bars: 15 μm.

The germ cell-to-maternal transition occurs during the cyst stages of oogenesis and is marked by the silencing of early oogenesis genes and upregulation of maternal genes (**Figure 1A1)** (Sarkar et al., 2023; DeLuca et al., 2020; Kotb et al., 2024; Blatt et al., 2021). Silencing of the early oogenesis genes *ribosomal small subunit protein 19b* (*rpS19b*) and *blanks* is frequently used as molecular readout of the germ cell-to-maternal transition (McCarthy et al., 2022; Sarkar et al., 2023). Despite the precise timing of this transition, how silencing of early oogenesis genes is coordinated with oocyte specification remains unclear.

The transcriptional silencing of early oogenesis genes requires the histone methyltransferase SET Domain Bifurcated Histone Lysine Methyltransferase 1 (SETDB1), which deposits Histone 3 Lysine 9 trimethylation (H3K9me3) to establish heterochromatin. (Clough et al., 2007; Rangan et al., 2011). Once silenced, these heterochromatic regions are demarcated by proteins such as Stonewall (Stwl) and anchored to the nuclear periphery via the nuclear pore complex (NPC) to maintain the silenced state (Sarkar et al., 2023; Kotb et al., 2024). Disruption of SETDB1, Stwl, or specific Nucleoporins (Nups) within the NPC results in aberrant expression of early oogenesis genes after the cyst stages, causing failed egg chamber growth and sterility (Maines et al., 2007; Clough et al., 2007; Sarkar et al., 2023; Kotb et al., 2024; Gigliotti et al., 1998). While these findings highlight the importance of chromatin reorganization in the germ cell-to-maternal transition, the upstream mechanisms driving this process remain unknown.

The Target of Rapamycin (TOR) pathway is a pivotal regulator of stem cell self-renewal and differentiation across many organ systems including Drosophila oogenesis (Kim and Sabatini, 2004; Wilson et al., 2024; Sanchez et al., 2016; Martin et al., 2022). TOR activity integrates nutrient availability and energy status to promote anabolic processes such as ribosome biogenesis, protein synthesis, and cell growth (Laplante and Sabatini, 2012; Johnson et al., 2013). For instance, TOR activity activation relieves the translational repression of terminal oligo-pyrimidine (TOP)-containing mRNAs to support efficient ribosome production that promotes GSC differentiation in Drosophila (Martin et al., 2022). TOR exists in two distinct complexes: Target of Rapamycin Complex 1 (TORC1) and TORC2, each with specific functions and downstream targets (Zoncu et al., 2011). TORC1 activity is dynamically regulated by GAP Activity Toward Rags 1(GATOR1) and GATOR2 during Drosophila oogenesis: GATOR1 promotes meiotic entry by inhibiting TORC1, whereas GATOR2 ensures TORC1 is reactivated in later stages to sustain differentiation. This balance between TORC1 inhibition and activation is essential for proper oogenesis. Loss of the GATOR2 components *missing oocyte* (*mio*) and *seh1* (also known as *Nup44A*) leads to arrested egg chamber growth and loss of oocyte fate (Iida and Lilly, 2004; Wei et al., 2014).

Intriguingly, a hypomorphic allele of the ribosomal protein gene *rpS2* (Cramton and Laski, 1994), depletion of Zinc Finger RP 8 (Zfrp8), a protein that affects cytoplasmic stability of ribosomal proteins including RpS2 (Minakhina et al., 2016), and loss of GATOR2 components *mio* and *Nup44A* phenocopy the loss of *SETDB1*, *stwl*, and Nups, leading to egg chambers that do not grow and a loss of oocyte fate (Iida and Lilly, 2004; Clough et al., 2007; Kotb et al., 2024; Maines et al., 2007; Gigliotti et al., 1998; Sarkar et al., 2023). Together, these findings raise the question of whether ribosome biogenesis and TORC1 activation orchestrate chromatin transitions across the cyst stage to ensure robust oocyte fate specification.

## Results

### Ribosome biogenesis, TORC1, and translation factors are required in the cyst stages for silencing *rpS19b* and *blanks*

To investigate the role of ribosome biogenesis in the germ cell-to-maternal transition, we depleted candidate genes that directly or indirectly regulate ribosome biogenesis and translation in the germline. These candidates included ribosome biogenesis regulators (*aramis*, *zfrp8*, *bystin*), TORC1 signaling components (*mio*, *raptor*) and translation elongation factor (*eEF1α1*) (**Figure 1B**) (Minakhina et al., 2016; Martin et al., 2022; Fukuda et al., 2008; Iida and Lilly, 2004; Wei et al., 2014). We used the germline-specific *nanos* GAL4 (*nos*GAL4) driver to induce RNA interference (RNAi) in the *rpS19b::GFP* reporter line under moderate knockdown conditions, preventing excessively severe depletion to avoid GSC and cyst differentiation defects (Doren et al., 1998; McCarthy et al., 2022; Martin et al., 2022). Ovaries from control and ribosome regulator germline knockdown (GKD) lines were stained for GFP and 1B1, the latter of which marks somatic cell membranes, spectrosomes, and fusomes in the germline (Zaccai and Lipshitz, 1996). We found that GKD of ribosome biogenesis regulators *aramis*, *zfrp8*, and *bystin*, TORC pathway members, *mio* and *raptor*, and the translation elongation factor *eEF1α1*, resulted in egg chambers that failed to grow (**Figure 1C-I1, Figure S1A**). In control ovaries, RpS19b::GFP expression was confined to undifferentiated stages and repressed in the egg chambers. In contrast, ovaries with GKD of ribosome biogenesis factors, TORC1 regulators, and the elongation factor exhibited aberrant expression of RpS19b::GFP in egg chambers (**Figure 1C-J, Figure S1B**). These findings suggest that regulators of ribosome biogenesis, TORC pathway members, and translation promote silencing of RpS19b::GFP in egg chambers and are required for egg chamber growth.

Early oogenesis genes are initially silenced in the cyst stages coinciding with elevated TORC activity (**Figure S1C-C1)**(McCarthy et al., 2022; Sarkar et al., 2023; Martin et al., 2022; Sanchez et al., 2016). To determine whether reducing ribosome levels or translation during this stage affects gene silencing, we used *bag of marbles-*GAL4 (*bam*GAL4), which is active in the cyst stages, to drive RNAi (Chen and McKearin, 2003). We stained ovaries from control and cysts-specific GKD lines of *zfrp8*, *mio*, *raptor*, and *eEF1α1* for Blanks, and 1B1 (Zaccai and Lipshitz, 1996). Compared to controls, the loss of *zfrp8*, *mio*, *raptor*, and *eEF1α1* in cyst stages resulted in egg chambers that failed to grow and exhibited persistent Blanks expression (**Figure S1D-H)**. These findings suggest that ribosome biogenesis, TORC1, and translation are required during the cyst stages to silence Blanks and promote egg chamber growth.

### Components that support translation are required for silencing early oogenesis genes

To determine whether ribosome-level regulators broadly influence gene expression during germ cell-to-maternal transition, we first performed RNA-sequencing (RNA-seq) on control and *zfrp8*, *mio*, and *raptor* GKD ovaries, as these genotypes yielded sufficient material for analysis. We used a 2-fold cut-off (Fold Change (FC)=2) and False discovery rate (FDR)<0.05) to identify dysregulated genes (**Figure 2A**). The overlapping set of genes in the three GKD ovaries reports on RNAs in the germline that are affected by the dysregulation of ribosome homeostasis. We detected 1483 genes that were upregulated in all three GKD ovaries (*zfrp8, mio* and *raptor*) compared to controls, constituting 50% of all upregulated genes (**Figure 2B, Supplementary Table 1**).

**Figure 2.**
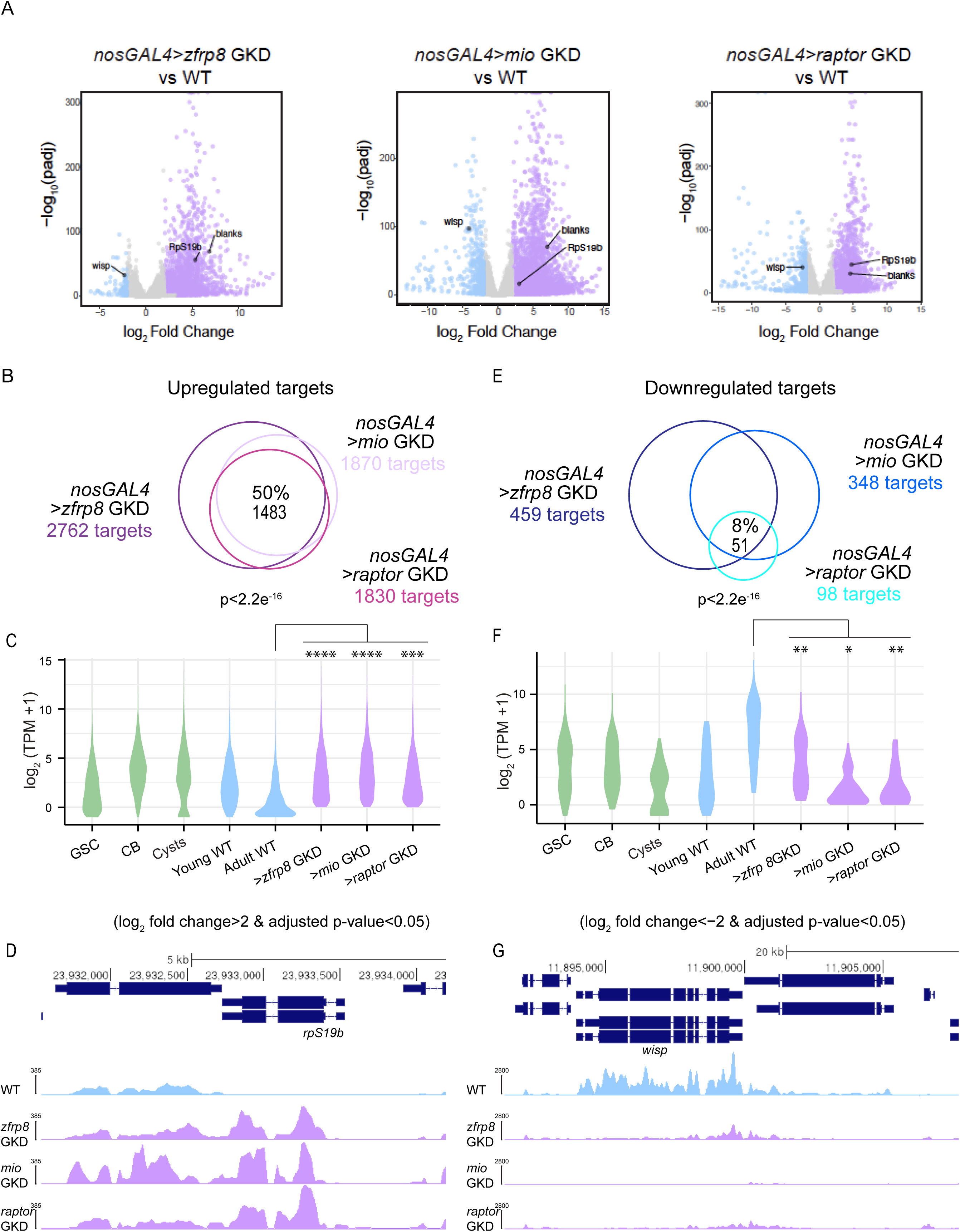
Ribosome level regulators are required for silencing early oogenesis genes during oogenesis. (A) Volcano plots of −log_10_ adjusted p value vs. log_2_ fold change (FC) of *nosGAL4* (control) vs >*zfrp8* GKD, > *mio*GKD and >*raptor* GKD ovaries showing significantly downregulated (blue) and upregulated (lilac) transcripts in *zfrp8* GKD, *mio* GKD and *raptor* GKD ovaries compared to control ovaries (FDR [false-discovery rate] < 0.05 and genes with 2-fold or higher change were considered significant). n=2 (B) Venn diagram of upregulated genes from RNA-seq of *>zfrp8* GKD, >*mio* GKD and >*raptor* GKD ovaries compared to controls. 50% of targets are shared upregulated targets, suggesting that ribosome level regulators are required to upregulate a cohort of genes during oogenesis. Statistics: Hypergeometric test p < 2.2e^-16^. (C) Violin plot of RNA levels of the shared upregulated targets between ribosome level regulators in ovaries enriched for GSCs, CBs, cysts, and whole ovaries, showing that the upregulated targets are expressed up to the cyst stages and attenuated in whole ovaries. Statistics: A negative binomial regression model was used to estimate the average TPM count of ‘signature’ genes between each ‘genotype’. The TPM of each gene was used as the dependent variable and ‘genotype’ was the independent variable. Statistical comparisons between groups were performed using contrasts and p-values were adjusted for multiple comparisons using the Benjamini-Hochberg procedure (FDR). The average TPM between groups was significantly different when p-FDR < 0.05. ns, p > 0.05; ∗p < 0.05; ∗∗p < 0.01; ∗∗∗p < 0.001; ∗∗∗∗p < 0.0001. (D) RPKM-normalized RNA-seq tracks showing that *rpS19b* is upregulated upon >*zfrp8* GKD, >*mio* GKD and >*raptor* GKD in dark purple compared to control (*nosGAL4)* (blue). (E) Venn diagram of downregulated genes from RNA-seq of *nosGAL4 > zfrp8* GKD ovaries *mio* GKD ovaries and *raptor* GKD ovaries compared to controls. 8% of the targets are shared downregulated targets. Statistics: Hypergeometric test p < 2.2e^-16^. (F) Violin plot of RNA levels of the shared downregulated targets between ribosome level regulators in ovaries enriched for GSCs, CBs, cysts, and whole ovaries, showing that the downregulated targets are expressed at higher levels in the differentiated stages. Statistics: A negative binomial regression model was used to estimate the average TPM count of ‘signature’ genes between each ‘genotype’. TPM of each gene was used as the dependent variable, and ‘genotype’ was the independent variable. Statistical comparisons between groups were performed using contrasts and p-values were adjusted for multiple comparisons using the Benjamini-Hochberg procedure (p-FDR). The average TPM between groups was significantly different when p-FDR < 0.05. ns, p > 0.05; ∗p < 0.05; ∗∗p < 0.01; ∗∗∗p < 0.001; ∗∗∗∗p < 0.0001. (G) RPKM-normalized RNA-seq tracks showing that *wisp* is downregulated upon >*zfrp8* GKD, >*mio* GKD and >*raptor* GKD in dark purple compared to control (*nosGAL4)* (blue).

To determine when these 1483 genes are typically expressed during development, we examined RNA-seq libraries enriched for specific stages of oogenesis, including undifferentiated stages (GSCs and CBs), differentiating stages during oocyte specification (cysts), and differentiated stages (early and late egg chambers in young and adult flies, respectively)(McCarthy et al., 2022; Blatt et al., 2021). We found that the 1483 genes are typically expressed in undifferentiated stages and are repressed as differentiation progresses, consistent with their classification as early oogenesis genes (**Figure 2C**). Importantly, these genes, including *rpS19b* and *blanks*, were expressed in late egg chambers from GKD of *zfrp8*, *mio*, or *raptor* but not WT controls (**Figure 2C-D, Figure S2A**). Thus, ribosome biogenesis and TORC1 activity is required to promote the silencing of early oogenesis genes at the onset of oocyte specification.

Additionally, we detected 51 genes that were downregulated in the *zfrp8, mio* and *raptor* GKD ovaries compared to controls, constituting 8% of all downregulated genes (**Figure 2E**). Using the stage-specific RNA-seq libraries, we found that these 51 genes are highly expressed in egg chambers of WT ovaries but not GKD of *zfrp8*, *mio*, or *raptor* egg chambers (**Figure 2F)**. Notably, these dysregulated genes encode maternally deposited transcripts, such as *wispy* and *zelda* (**Figure 2G**, **Figure S2B)**. Therefore, ribosome biogenesis and TORC1 activity are also essential for upregulating a small subset of genes, including those that are required to launch the next generation.

### Ribosome biogenesis, TORC1 activity, and heterochromatin machinery converge on silencing of early oogenesis genes

The H3K9me3 histone methyltransferase, *SETDB1,* genome organization protein such as *Nup154* (a component of the NPC), and the H3K27me3 histone methyltransferase *enhancer of zeste* (*e(z)*) are required for silencing of early oogenesis genes in differentiated egg chambers and for egg chamber growth (Sarkar et al., 2023; DeLuca et al., 2020). The phenotypic similarity between ribosome-level regulators and chromatin modulators suggested a potential link between ribosome biogenesis, TORC1, and chromatin regulation during oocyte development. To test this, we plotted log_2_FC of genes in *zfrp8*, *mio*, and *raptor* GKD with previously published datasets for *SETDB1*, *Nup154*, and *e(z)* GKD ovaries(Sarkar et al., 2023; DeLuca et al., 2020). Strikingly, 47% (658 genes) of the upregulated targets in ribosome regulators *zfrp8*, *mio*, and *raptor* GKD overlapped with those in *SETDB1*, *Nup154*, and *e(z)* GKD (**Figure 3A, Supplementary Table 2**). These genes, which are typically repressed in differentiated stages, are associated with quiescent chromatin (**Figure 3A**). This overlap suggests that translation regulators and chromatin modulators co-regulate the silencing of a shared set of early oogenesis genes.

**Figure 3.**
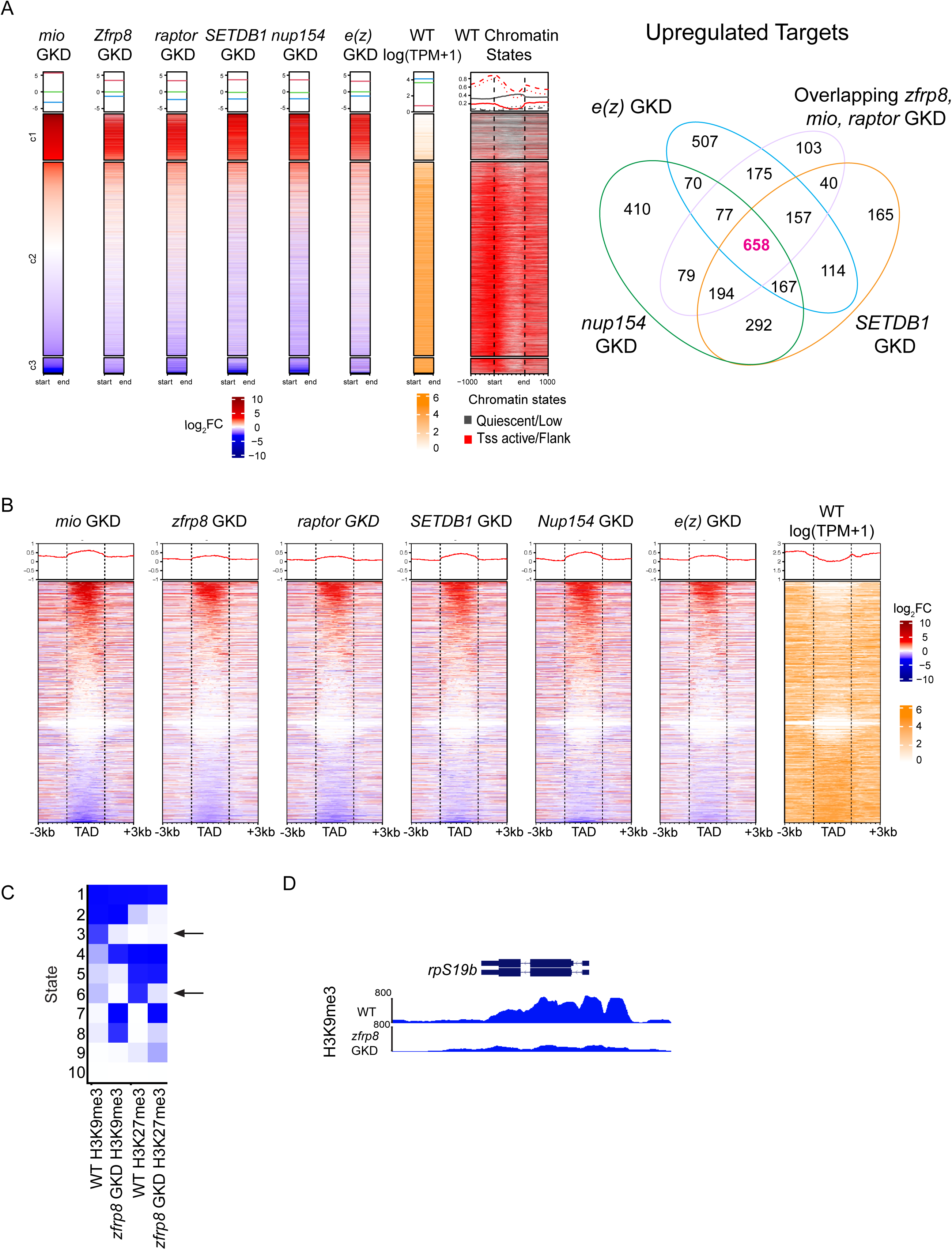
Ribosome level regulators control a cohort of common targets with heterochromatin modulators. (A) Heatmaps of log_2_FC sorted genes *zfrp8* GKD, *mio* GKD and *raptor* GKD along with *SETDB1* GKD, *Nup154* GKD and *e(z)* GKD targets showing they all share a common cohort of upregulated and downregulated targets. Upregulated targets are shown in red and downregulated targets are shown in blue. Heatmaps show chromatin states for those targets in differentiated stages of wild-type ovaries. Venn diagram showing overlapping upregulated targets between *zfrp8* GKD, *mio* GKD and *raptor* GKD with *SETDB1* GKD, *e(z)* GKD and *Nup154* GKD. The overlap is 658 genes (47%) showing that ribosome level regulators and heterochromatin regulators share a common cohort of upregulated targets during oogenesis. Statistics: Hypergeometric test p < 2.2e^-16^ (B) Heat maps of log_2_FC sorted for RNA-seq targets of ribosome regulators at TADs. Upregulated targets are shown in red, and downregulated targets are shown in blue. Heat maps are shown for TADS as well as −3 kb and +3 kb around the start and end of TADs. Black dashed lines represent the start and end of TADs. The heat maps are divided into three clusters: the upregulated target cluster, the nontarget gene cluster, and the downregulated target cluster. The heat maps show that both genes upregulated and genes downregulated upon ribosome regulator GKD are present within TADs. (C) A chromatin model for H3K9me3 and H3K27me3 regulation in WT and >*zfrp8* GKD ovaries. In *zfrp8* GKD, H3K9me3 levels are reduced in state 3 while H3K27me3 is redistributed, indicating chromatin instability. These findings suggest that ribosome biogenesis is required for maintaining epigenetic silencing during oocyte specification. (D) Tracks showing the level of H3K9me3 on *rps19b* locus in control and *zfrps8* GKD showing reduction of heterochromatin.

Our previous findings revealed that certain early oogenesis genes are not directly regulated by heterochromatin but rather become misexpressed due to their localization within topologically associating domains (TADs) (Kotb et al., 2024). Genes positioned within these domains appear especially vulnerable to global shifts in chromatin modifications or changes in nuclear architecture (Jerković et al., 2020; Kotb et al., 2024). For instance, while the gene *blanks* is not directly targeted by the heterochromatin-depositing enzyme SETDB1, its silencing depends on intact genome architecture mediated by NPCs (Kotb et al., 2024; Sarkar et al., 2023). In contrast, *rpS19b* carries heterochromatic marks on its promoter and gene body (Kotb et al., 2024; Sarkar et al., 2023).

Our previous work demonstrated that disrupting heterochromatin or nuclear pore complexes (NPCs)—which are interlinked through a feedback loop—leads to compromised genome architecture and a failure to silence these early oogenesis genes (Sarkar et al., 2023)(Kotb et al., 2024). To further explore this connection, we mapped the genes dysregulated upon loss of ribosome-level regulators in relation to TADs. This analysis revealed that these genes predominantly reside within TADs and are silenced in differentiated stages (**Figure 3B**).

Given their co-regulation of target genes and their similar phenotypes, we hypothesized that *zfrp8*, *mio*, and *raptor* GKD could perturb chromatin state and genome organization, similar to GKD of *SETDB1*, *Nup154* and *e(z).* To test this, we performed Cleavage Under Targets and Release Using Nuclease (CUT&RUN) assays to assess levels of the H3K9me3 and H3K27me3 chromatin marks—deposited by SETDB1 and E(z), respectively—in control and *zfrp8* GKD ovaries (Skene and Henikoff, 2017). We analyzed H3K9me3 and H3K27me3 chromatin marks in WT and *zfrp8* GKD ovaries to define 10 genome-wide states based on their distribution patterns. Our analysis revealed a significant decrease in both H3K9me3 and H3K27me3 levels on a subset of genes in *zfrp8* GKD ovaries (**Figure 3C**, **Supplementary Table 3**). However, we also observed a redistribution of these marks, with their loss at specific loci accompanied by their gain elsewhere in the genome (**Figure 3C**). Notably, the loss of H3K9me3 was particularly evident at genes with promoter-associated H3K9me3 (**Figure S3A-B**). Among these, *rpS19b* exhibited a striking reduction in H3K9me3 levels, consistent with its transcriptional upregulation in *zfrp8* GKD ovaries (**Figure 3D**). These findings suggest that maintaining adequate ribosome level regulators is essential for proper chromatin state either directly or indirectly contributing to the silencing of early oogenesis genes during oocyte specification.

### Ribosome biogenesis promotes translation of the nucleoporin and TORC1 regulator Nup44A

As we found that proper ribosome levels, TORC1 activity and translation are crucial to silence early oogenesis genes, we sought to identify the targets that could be translationally affected when ribosome level regulators are depleted. To investigate this, we performed a polysome-seq analysis on ovaries from *zfrp8* GKD and control flies (**Figure 4A**) (McCarthy et al., 2022)(Breznak et al., 2023). We observed a reduction in the levels of the small (40S) and large (60S) ribosomal subunits, and monosomes (80S), in *zfrp8* GKD compared to WT, consistent with its role in ribosome biogenesis (**Figure 4B**) (Breznak et al., 2023). By comparing polysome-associated RNAs to total RNAs, we identified mRNAs whose translation was disrupted in *zfrp8*GKD ovaries relative to wild-type controls (**Figure 4C**)(Ernlund et al., 2018). Our analysis revealed that 536 transcripts were translationally downregulated and 509 were translationally upregulated in *zfrp8* GKD ovaries indicating widespread translational dysregulation.

**Figure 4.**
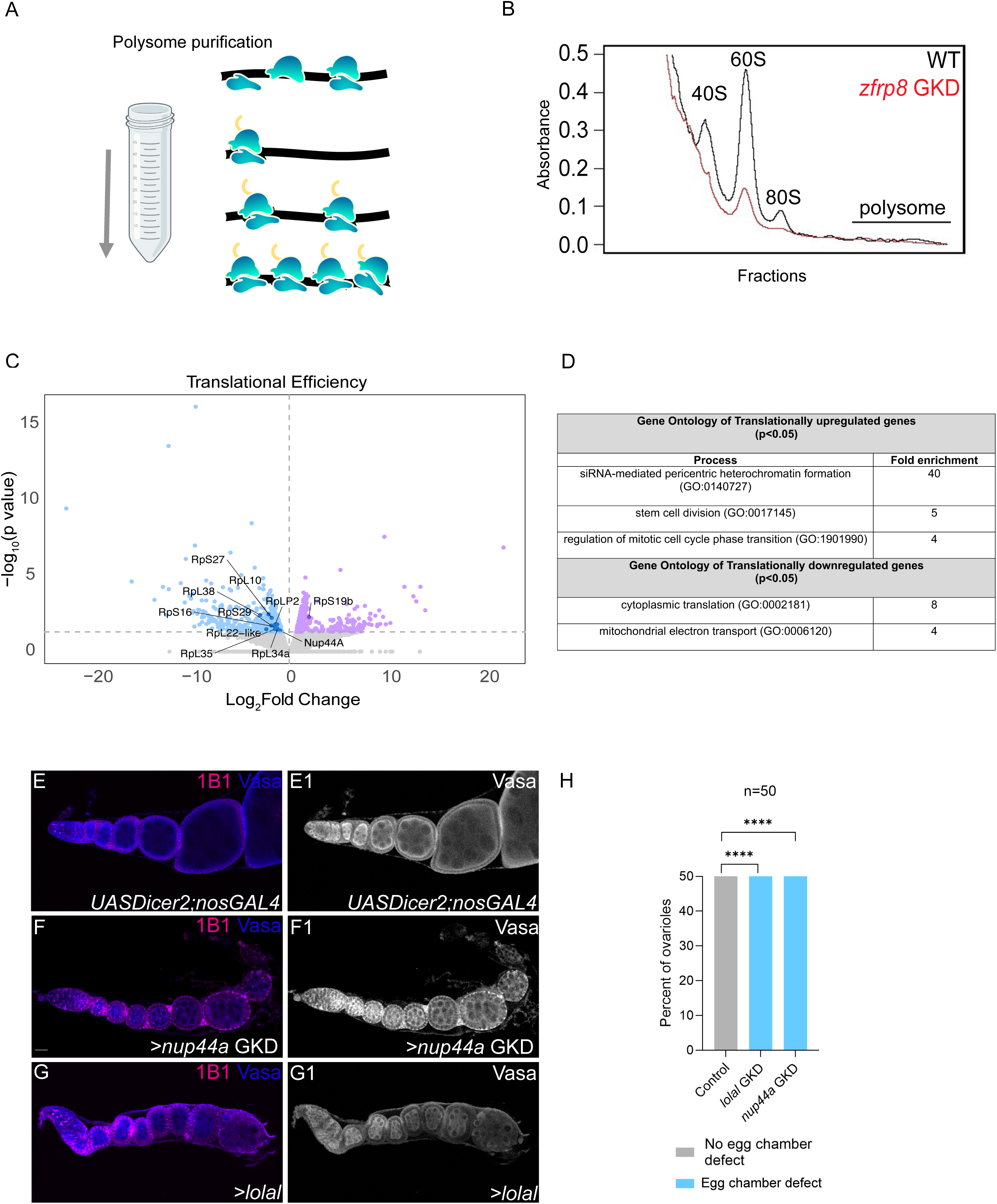
Adequate ribosome levels are required for the translation of an NPC component. (A) A schematic of the polysome isolation from control *nosGAL4* and *>zfrp8* GKD ovaries using sucrose gradients. (B) Graph depicting 40S, 60S subunits, monosomes and polysomes isolated from control *nosGAL4* and >*zfrp8* GKD ovaries. *zfrp8* is required in the germline for proper formation of 40S, 60S and monosomes. (C) Graph showing the translation efficiency comparing translational targets of >*zfrp8* GKD ovaries compared to *nosGAL4* (control) varies. Downregulated translational targets upon depletion of *zfrp8* are shown in blue, and upregulated in lilac. (D) Gene ontology of translationally upregulated and downregulated targets. (E–E1) Ovariole of control *rpS19b::GFP;nosGAL4* (D) and grayscale (D1) stained for GFP (green) and 1B1 (magenta). In control, GFP is expressed in the undifferentiated stages and early cysts and then silenced. (E-E1) Ovariole of *Nup44A* GKD (E) and grayscale (E1) stained for Vasa (green, right grayscale) and 1B1 (magenta). *Nup44A* GKD resulted in egg chambers did not grow and died during oogenesis. (F) Ovariole of *lolal* GKD (E) and grayscale (E1) stained for Vasa (green, right grayscale) and 1B1 (magenta). *lolal* GKD resulted in egg chambers that did not grow and died during oogenesis. (H) Quantitation of percent ovarioles with lack of egg chamber growth phenotype in *Nup44A* GKD and *lolal* GKD, compared to control ovaries. Statistics: Two-tailed t-test; n = 50 ovarioles per genotype; ns, p > 0.05; ∗p < 0.05; ∗∗p < 0.01; ∗∗∗p < 0.001; ∗∗∗∗p < 0.0001.

Gene ontology (GO) analysis of the upregulated targets, included proteins involved in siRNA-mediated pericentric heterochromatin and proteins associated with chromatin organization, such as Suppressor of hairy wing (Su(Hw)) and M1BP, suggesting potential feedback regulation triggered by the loss of heterochromatin (**Figure 4C-D, Supplementary Table 3**)(Bag et al., 2021)(Baxley et al., 2011). In addition, genes involved in stem cell division were upregulated indicating an impaired transition from a GSC-like program to an oocyte program.

Conversely, GO analysis of translationally downregulated transcripts revealed a significant reduction in ribosomal components (**Figure 4D)**, consistent with previous reports that loss of ribosome biogenesis components decreases ribosomal protein synthesis (Martin et al., 2022; Breznak et al., 2023). However, *rpS19b* was not among the downregulated ribosomal proteins, consistent with the observation that it is ectopically expressed upon loss of *zfrp8*. Additionally, visual inspection of the downregulated targets identified genes such as *Nup44A*, which encodes a nuclear pore protein and *lola-like*/*batman*, which belongs to a family of BTB/POZ domain and interacts with both Polycomb and Trithorax complexes (**Supplementary Table 3**)(Senger et al., 2011; Faucheux et al., 2003). We found that independent GKD of either *Nup44A* or *lola-like*/*batman* resulted in egg chambers that do not grow and sterility phenocopying loss of ribosome regulators and heterochromatin components (**Figure 4E-H)**. Thus, depletion of *zfrp8* results in translational dysregulation of components that regulate either proper chromatin state or genome architecture during oocyte specification.

### Nup44A is critical for silencing early oogenesis genes

Seh1, the mammalian homolog of Drosophila Nup44A, is a multifunctional protein essential for NPC assembly and TORC1 signaling regulation (von Appen et al., 2015; Senger et al., 2011). Within the NPC, Nup44A is a core component of the Nup107-160 subcomplex, a structural scaffold critical for NPC assembly, nuclear-cytoplasmic transport, and chromatin organization (Walther et al., 2003). This complex is particularly important during development, where it supports chromatin interactions and epigenetic regulation (Gozalo et al., 2020). Loss of Nup44A disrupts nuclear basket assembly (Senger et al., 2011). Beyond its structural role in the NPC, Nup44A also modulates TORC1 signaling, integrating nuclear transport with metabolic regulation (Senger et al., 2011). This dual function positions Nup44A as a crucial link between nuclear transport, metabolic signaling, and epigenetic regulation during development. Given its central role, we focused on Nup44A for further analysis.

In Drosophila, *Nup44A* mutants are homozygous viable but largely sterile due to a loss of oocyte specification (Senger et al., 2011). Since Zfrp8 regulates early oogenesis genes and is required for Nup44A translation, we hypothesized that Nup44A is similarly essential for silencing early oogenesis genes. To test this, we depleted *Nup44A* in the germline and assessed egg chamber growth and the expression of the *rpS19b::GFP* reporter. We found that GKD of *Nup44A* resulted in RpS19b::GFP expression in differentiated egg chambers and led to growth-arrested egg chambers, phenocopying the loss of heterochromatin components and ribosome regulators (**Figure 5A-C**). To further investigate whether *Nup44A* regulates early oogenesis genes, we performed RNA-seq on *Nup44A* GKD ovaries, comparing gene expression profiles with those of ribosome regulators *zfrp8*, *mio*, *raptor,* and heterochromatin regulators *SETDB, Nup154,* and *e(z)*. We identified 2375 upregulated genes (**Figure 5D-E, Supplementary Table 4**). Out of the 658 genes upregulated in *zfrp8*, *mio*, *raptor*, *SETDB1*, *Nup154*, and *e(z)* GKD, 97% are also upregulated in *Nup44A* GKD (**Figure 5E, Supplementary Table 4**). Notably, genes such as *rpS19b*, which normally carry promoter-associated H3K9me3, showed a reduction in this mark, consistent with their transcriptional upregulation in *Nup44A* GKD ovaries (**Figure 5D-E**). These findings establish Nup44A as a key regulator of early oogenesis gene silencing, linking NPC function to ribosome-mediated translational control during oocyte specification.

**Figure 5:**
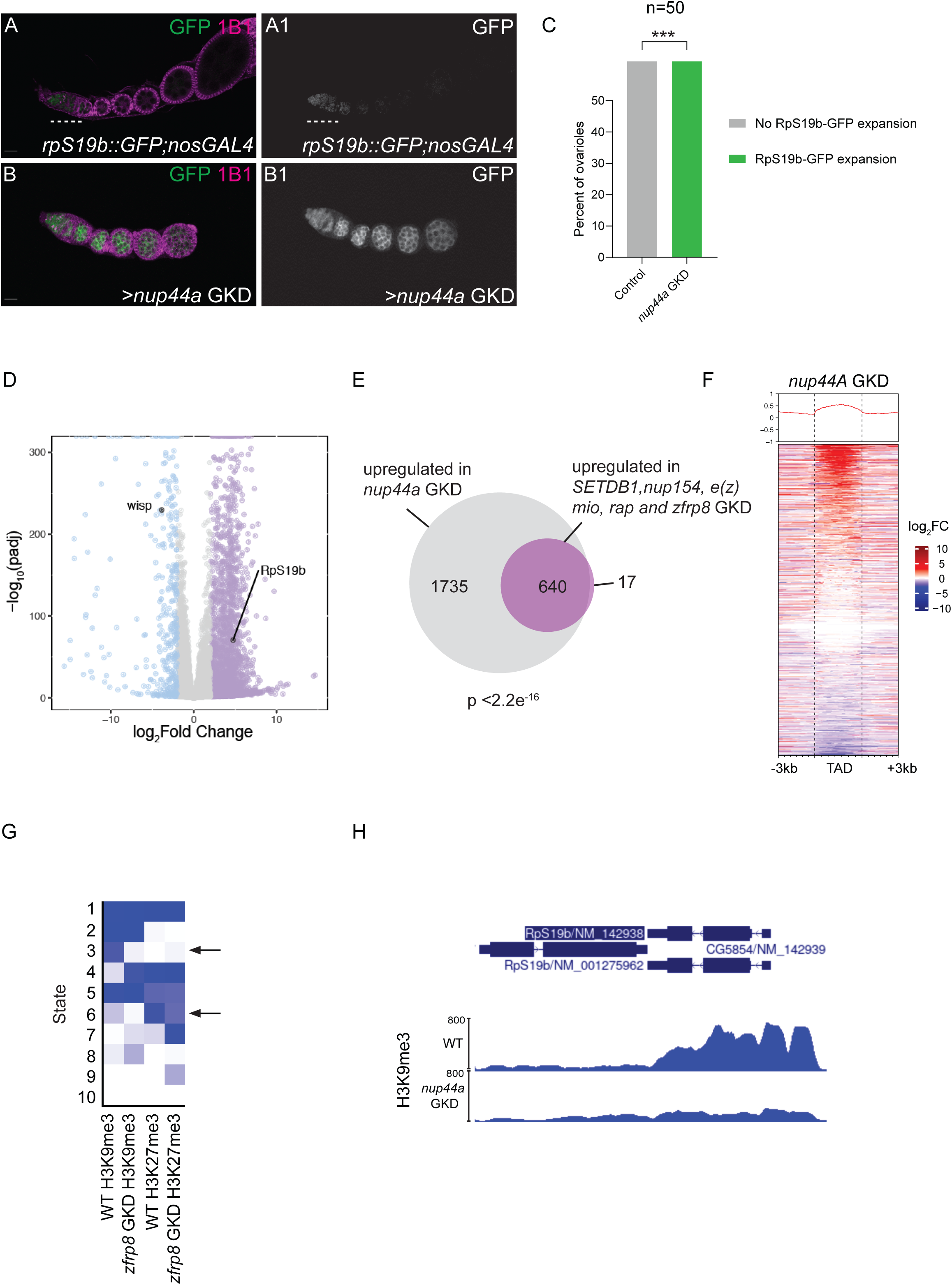
Nup44A is required for silencing early oogenesis genes and promoting a stable epigenetic state. (A–A1) Ovariole of control *rpS19b::GFP;nosGAL4* (A) and grayscale (A1) stained for GFP (green) and 1B1 (magenta). In control, GFP is expressed in the undifferentiated stages and early cysts and then silenced. (B-B1) Ovariole of *Nup44A* GKD (B) and grayscale (B1) stained for GFP (green, right grayscale) and 1B1 (magenta). *Nup44A* GKD resulted in egg chambers that ectopically expressed RpS19b::GFP (white dashed line), did not grow, and died during oogenesis. (A) Quantitation of percent ovarioles with RpS19b::GFP expression. Statistics: Two-tailed t-test; n = 50 ovarioles per genotype; ns, p > 0.05; ∗p < 0.05; ∗∗p < 0.01; ∗∗∗p < 0.001; ∗∗∗∗p < 0.0001. (B) Volcano plots of −log_10_ adjusted p value vs. log_2_fold change (FC) of *nosGAL4*> *nup44a* GKD ovaries vs *nosGAL4* (control) showing significantly downregulated (blue) and upregulated (lilac) genes in *nup44a* GKD ovaries (FDR < 0.05 and genes with 2-fold or higher change were considered significant). n=2 (C) Venn diagram of upregulated genes from RNA-seq of >Nup44A compared to ribosome level and chromatin regulators in Figure 3. Statistics: Hypergeometric test p < 2.2e^-16^. (D) RPKM-normalized tracks showing the level of H3K9me3 on *rps19b* locus in control and *Nup44A* GKD showing reduction of heterochromatin. (E) A chromatin model for H3K9me3 and H3K27me3 regulation in WT and >*Nup44A* GKD ovaries. In *Nup44A* GKD, H3K9me3 levels are reduced on promoters in state 3. (F) RPKM-normalized tracks showing the level of H3K9me3 on *rps19b* locus in control and *Nup44A* GKD showing reduction of heterochromatin.

Previous studies had shown that NPC components are frequently enriched at TAD boundaries and that a large fraction of early oogenesis genes reside within these domains (Kotb et al., 2024). We also found that genes dysregulated by ribosome-level regulators are predominantly located within TADs (**Figure 3B)**. Building on this, we asked whether genes affected by *Nup44A* depletion were also situated within TADs. By mapping genes dysregulated upon loss of *Nup44A*, we observed that these early oogenesis genes, much like those regulated by SETDB1 and the NPC component Nup154, are located within TADs (**Figure 5F)**. To investigate whether *Nup44A* regulates chromatin state, we performed CUT&RUN analysis for the heterochromatin marks H3K27me3 and H3K9me3 in *Nup44A* GKD ovaries. Similar to *zfrp8* depletion, we observed significant alterations in H3K27me3 and H3K9me3 levels, with their loss at specific loci accompanied by their gain elsewhere in the genome (**Figure 5G, Figure S5A-B**). Notably, the loss of H3K9me3 was particularly evident at genes with promoter-associated H3K9me3 on genes typically enriched for these marks such as *rpS19b* (**Figure 5H)**. These findings reinforce the role of *Nup44A* in maintaining proper chromatin state and genome architecture, further supporting the model that ribosome biogenesis and NPC integrity work together to establish the epigenetic landscape necessary for oocyte specification.

### Nup44A supports a feedforward loop between TORC1 signaling and ribosome biogenesis to drive oogenesis

We next sought to determine how *Nup44A* translation is regulated by ribosome levels. We previously observed that terminal oligo-pyrimidine (TOP)-containing RNAs are translationally downregulated upon reduction of ribosome biogenesis in the GSCs (Martin et al., 2022). These motifs are known to coordinate the translation of specific proteins in response to TORC1 activity (Thoreen et al., 2012; Lahr et al., 2017; Fonseca et al., 2018; Philippe et al., 2018). Indeed, upon GKD of *zfrp8,* we observed that several ribosomal proteins are translationally downregulated (**Figure 4C-D)**. We hypothesized that the downregulation of *Nup44A* mRNA translation could be mediated by a potential TOP motif within its 5′ untranslated region (5′ UTR). To test this, we first asked if *Nup44A* 5′ UTR contained a TOP motif. Cap analysis of gene expression sequencing (CAGE-seq) data from total mRNA from the ovary was used to determine transcription start sites (TSSs) to accurately determine the 5’ end of transcripts. TOP sequences start with a C or U base and contain a run of pyrimidine bases (Chen et al., 2014). We found that *Nup44A*-RA transcript was transcribed during oogenesis and the 5’UTR initiated with a C base and contained a stretch of pyrimidines (**Figure 6A)**. Thus, *Nup44A* is a TOP containing mRNA.

**Figure 6.**
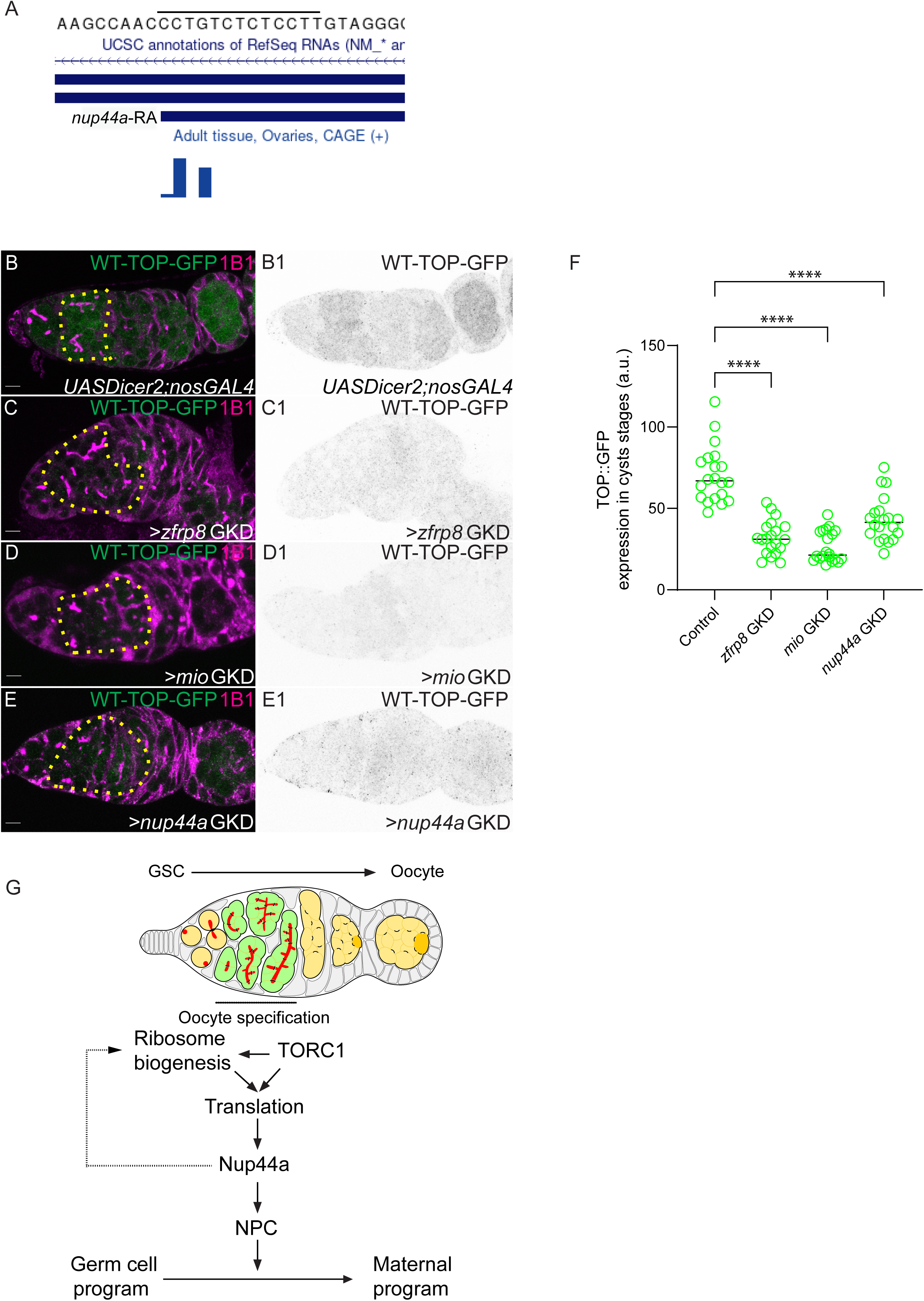
*Nup44A* contains a TOP-motif and TOP-mRNA translation is sensitive to ribosome level regulators in the cyst stages. (A) CAGE-Seq analysis of the *Nup44A* transcript start site, showing an enrichment of pyrimidines, consistent with a terminal oligo-pyrimidine (TOP) motif, which is a feature of TORC1-sensitive mRNAs. (B-B1) Germarium from a TOP-reporter::GFP control stained for GFP (green, right in grayscale) and 1B1 (magenta). TOP reporter expression is observed in the cyst stages and persists into the first egg chamber. (C-C1) Germarium from TOP-reporter::GFP >*zfrp8* GKD stained for GFP (green, right in grayscale) and 1B1(magenta). TOP reporter expression is attenuated in the cyst stages upon >*zfrp8* depletion compared to control. (D-D1) Germarium from TOP-reporter::GFP >*mio* GKD stained for GFP (green, right in grayscale) and 1B1(magenta). TOP reporter expression is attenuated in the cyst stages upon >*mio* depletion compared to control. (E-E1) Germarium from TOP-reporter::GFP >*Nup44A* GKD stained for GFP (green, right in grayscale) and 1B1(magenta). TOP reporter expression is attenuated in the cyst stages upon >*Nup44A* depletion compared to control. (F) Quantification of GFP levels in the cyst stages of >*zfrp8* GKD, >*mio* GKD, and >*Nup44A* GKD compared to control ovaries. Loss of these factors results in reduced TOP reporter expression. Statistics: Two-tailed t-test; n = 50 ovarioles per genotype; ns, p > 0.05; *p < 0.05; **p < 0.01; ***p < 0.001; **p < 0.0001. (G) Model illustrating how TORC1-driven translation and ribosome biogenesis regulate heterochromatin establishment during oocyte specification. In Drosophila, Zfrp8 promotes the translation of Nup44A, a nuclear pore complex (NPC) component, which is required for proper chromatin organization. H3K9me3 marks promoters of the early oogenesis genes to maintain their repression, while H3K27me3 regulates developmental genes. Loss of Zfrp8, TORC1 activity, or NPC components disrupts translation, leading to chromatin instability and gene misregulation, ultimately affecting oocyte fate commitment. This translation-epigenetic axis ensures the irreversible transition from germ cell to oocyte. Scale bars: 7.5 μm.

To investigate whether the TOP motif contributes to the regulation of translation *in vivo* during the cyst stages, we crossed a previously characterized TOP reporter fly with *zfrp8* and *mio* GKD and monitored its translation levels in the cyst stages. Under normal conditions, the TOP reporter was strongly expressed in 8-cell cysts, reflecting its dependence on TORC1 activity (**Figure 6B-B1)**. However, upon GKD of *zfrp8* or *mio*, GFP expression from the TOP reporter was significantly reduced, suggesting that both *zfrp8* and *mio* are required for the efficient translation of TOP motif-containing RNAs (**Figure 6C-D1, F)**. As *Nup44A* is also implicated in the TORC1 pathway (Senger et al., 2011), we asked if GKD of *Nup44A* would also result in loss of TOP translation. Similar to the results with *zfrp8* and *mio* GKD, we found that *Nup44A* GKD led to a marked downregulation of GFP expression from the TOP reporter, further supporting the idea that ribosome level regulators mediate the translation of TOP motif-containing RNAs (**Figure 6C-F)**. Overall, our data reveal that translation of the TOP-containing *Nup44A* mRNA is reduced when ribosome biogenesis is impaired. Since Nup44A supports TORC1 signaling, which in turn drives ribosome biogenesis, its reduced translation weakens TORC1 activity, reinforcing a feedforward loop that further limits ribosome production.

## Discussion

### Translation regulates epigenetic reprogramming during oocyte specification

During oogenesis, germ cells undergo a tightly regulated differentiation process to give rise to a specified oocyte, which will mature into an egg capable of launching the next generation (Spradling et al., 2011; Lehmann, 2012). This process, termed germ cell-to-maternal transition, requires the silencing of early oogenesis genes and the activation of maternal genes, ensuring a stable and irreversible cell fate commitment (Sarkar et al., 2023; Kotb et al., 2024; Blatt et al., 2021). Here, we demonstrate that translation, supported by ribosome biogenesis and TORC1 activity, plays a pivotal role in regulating this epigenetic reprogramming. We identified ribosome biogenesis factors (Zfrp8, Bystin), TORC1 components (Raptor, Mio), and translation-related factors such as eEF1α1 as critical regulators of gene silencing during oogenesis. Our findings suggest that the timing of the germ cell-to-maternal transition is mediated by TORC1 activity, which increases during oocyte specification to promote translation of key factors required for chromatin remodeling and gene silencing. This regulatory mechanism ensures that early oogenesis genes are repressed precisely as maternal programs are activated, establishing an irreversible commitment to oocyte fate.

Mechanistically, our findings suggest that ribosome biogenesis and TORC1 activity regulate nuclear architecture by influencing the translation of key NPC components, such as Nup44A (Senger et al., 2011; Gozalo et al., 2020), as well as chromatin through the regulation of the transcription factor Lolal, which interacts with chromatin modifiers (Faucheux et al., 2003). Since NPC assembly depends on the coordinated expression of its components, perturbations in ribosome biogenesis or TORC1 activity disrupt proper NPC formation, leading to defects in heterochromatin maintenance (**Figure 6G**). Notably, our CUT&RUN analysis revealed that while H3K9me3 and H3K27me3 marks were reduced at certain loci, they were redistributed elsewhere in the genome. This suggests that most gene expression misregulation arises not solely from heterochromatin loss but from broader disruptions in genome organization. These findings support a model in which transient translational regulation establishes stable epigenetic states by promoting proper genome organization, ensuring irreversible cell fate transitions. By uncovering this translation-epigenetic axis, we reveal a previously unrecognized mechanism by which short-term fluctuations in translation are epigenetically encoded to drive long-term developmental decisions.

### Ribosome biogenesis orchestrates sequential transitions during germ cell differentiation into an oocyte

Ribosome biogenesis is a fundamental determinant of stem cell fate, balancing self-renewal and differentiation by regulating protein synthesis (Zhang et al., 2014; Sanchez et al., 2016; Khajuria et al., 2018). In many stem cell systems, high ribosome biogenesis sustains proliferation and maintains the undifferentiated state by coupling translational control with cell cycle progression. However, as stem cells commit to differentiation, ribosome biogenesis is often downregulated, leading to selective translation of lineage-specific factors that drive cell fate transitions (Khajuria et al., 2018; Sanchez et al., 2016). This shift ensures that differentiation-associated gene programs are properly executed while preventing premature or inappropriate cell fate decisions.

We previously found that ribosome biogenesis plays a crucial role in orchestrating the stepwise differentiation of GSCs into oocytes (Martin et al., 2022). In GSCs, high ribosome biogenesis supports cell cycle progression by regulating factors that suppress P53, enabling continued cell division (Martin et al., 2022). However, upon GSC division, ribosome biogenesis levels decrease, coinciding with an increase in translation that drives the expression of meiotic genes and promotes early differentiation (Breznak et al., 2023). As germ cells progress into meiosis, translation levels increase within the 8-cell cyst, where it becomes essential for the synthesis of key factors, including Nup44A and chromatin regulators such as Lolal. These findings highlight a dynamic interplay between ribosome biogenesis and translation, where transient changes in ribosome levels dictate sequential waves of translation that activate differentiation programs in a controlled, stepwise manner.

While ribosome levels and translation dictate which mRNAs are translated efficiently, additional layers of regulation fine-tune protein synthesis during differentiation. We propose that trans-acting factors such as LARP, Pumilio, and Bruno act as RNA-binding proteins that license specific mRNAs for translation at distinct developmental stages (Flora et al., 2018b; Blatt et al., 2020). Thus, a combination of ribosome biogenesis, TORC1 activity, and translational control mechanisms ensures the timely expression of critical differentiation factors, allowing for a regulated transition from GSC self-renewal to oocyte specification.

### TORC1 integrates translational and epigenetic control to ensure oocyte fate commitment

TORC1 is a central regulator of cell growth and metabolism, integrating nutrient availability and cellular energy status to control translation (Laplante and Sabatini, 2012; Kim and Sabatini, 2004). In stem cell differentiation, TORC1 activity must be precisely modulated to balance self-renewal with lineage commitment (Iida and Lilly, 2004; Wei et al., 2014). In the germline, TORC1 activation promotes translation of factors required for cell cycle progression and early differentiation. However, our findings reveal that TORC1’s role extends beyond merely increasing protein synthesis—it also influences epigenetic reprogramming by regulating the translation of chromatin modifiers and nuclear pore components. Through this mechanism, TORC1 establishes the chromatin landscape necessary for differentiation, ensuring the timely repression of germline genes and activation of oocyte-specific programs.

Depletion of ribosome biogenesis and TORC1 factors leads to the upregulation of early oogenesis genes and downregulation of maternal genes, phenocopying the loss of *SETDB1*, *e(z)*, and *Nup154*—key regulators of heterochromatin formation. This direct link between translation and chromatin regulation suggests that adequate ribosome levels are required for proper heterochromatin establishment during oocyte specification. Supporting this, we observe significant changes in H3K27me3 and H3K9me3 marks upon depletion of *zfrp8*, reinforcing the idea that ribosome biogenesis is essential for chromatin remodeling during differentiation. However, as noted earlier, redistribution of chromatin marks suggests that the primary defect may lie in genome organization rather than a simple loss of heterochromatin (Iida and Lilly, 2004).

### Nup44A links NPC assembly and TORC1 activity to coordinate translational and epigenetic regulation

Nup44A, a component of the NPC, plays a crucial role in coordinating translational control and epigenetic reprogramming during oocyte specification (Senger et al., 2011). NPCs regulate nucleocytoplasmic transport and are essential for establishing chromatin organization, particularly through interactions with heterochromatin (Gozalo et al., 2020; Sarkar et al., 2023; Capelson et al., 2010). Our findings indicate that *Nup44A* translation is tightly regulated by ribosome biogenesis and TORC1 activity, positioning it as a key factor that synchronizes these two processes. TORC1-driven translation of *Nup44A* ensures proper NPC function, which in turn facilitates heterochromatin formation required for gene silencing. Loss of *Nup44A* disrupts nuclear architecture, leading to defects in the deposition of H3K9me3 and H3K27me3 marks, phenocopying the effects of TORC1 and ribosome biogenesis depletion. This suggests that NPC integrity is not only structurally important but also functionally critical for maintaining the chromatin state necessary for irreversible cell fate transitions. By integrating translational control with nuclear organization, *Nup44A* serves as a molecular link that synchronizes the transient regulation of translation with the long-term establishment of the oocyte epigenome, ensuring the proper execution of germ cell-to-maternal transition.

Our findings, together with previous work, highlight the conserved role of nutrient-sensing pathways in orchestrating chromatin architecture to regulate cell fate decisions. While fasting-induced chromatin reorganization in *C. elegans* is mediated through mTOR inhibition and RNA Pol I (Al-Refaie et al., 2024), we show that in Drosophila, TORC1 activation during oocyte specification drives a translation-dependent mechanism that establishes stable heterochromatin states. This suggests that metabolic inputs dynamically regulate chromatin architecture, either by remodeling genome organization in response to environmental cues, as seen in fasting (Al-Refaie et al., 2024), or by ensuring the irreversible silencing of developmental programs during differentiation, as observed in oogenesis. A key link between these processes may be the nucleolus and NPC, both of which are remodeled in response to translation and mTOR activity (Al-Refaie et al., 2024).

Overall, our work uncovers how transient translational changes impact long-term chromatin states, suggesting that ribosome biogenesis and translation could serve as central regulators of epigenetic memory across diverse biological contexts. This raises intriguing possibilities that metabolic shifts could influence chromatin states not just in development but also in disease and aging, opening new avenues for exploring how translation-mediated chromatin regulation shapes cellular identity and adaptation (Al-Refaie et al., 2024). Thus, by elucidating this intricate interplay, we provide a foundation for future studies on how translation-mediated chromatin regulation contributes to developmental robustness and cellular plasticity.

## Materials and Methods

### Drosophila Stocks

This investigation utilized the following RNAi and mutant Drosophila lines: *zfrp8* RNAi (Bloomington #36581), *mio* RNAi (Bloomington #57745), *raptor* RNAi (Bloomington # 34814) *bystin* RNAi (Bloomington # 34876) and *aramis* RNAi (Bloomington #32334). The study also employed the RpS19b::GFP tagged line (McCarthy et al., 2022). Germline-specific drivers and double balancer lines used included: UAS-Dcr2;*nos*GAL4 (Bloomington #25751), *bam*GAL4 (Bloomington #80579), matGAL4 (Bloomington #7062,7063), *nos*GAL4;MKRS/TM6 (Bloomington #4442).

### Drosophila Husbandry

Fly crosses were cultivated at 25-29°C, with dissections performed between 0-3 days post-eclosion. Fly food for stocks and crosses was prepared following the previously published laboratory protocol (summer/winter mix). Narrow vials (Fisherbrand Drosophila Vials; Fisher Scientific) were filled to approximately 10-12mL(Flora et al., 2018a)(Upadhyay et al., 2018).

### Dissection and Immunostaining

Ovaries were extracted and ovarioles separated using mounting needles in PBS solution on ice. Samples were fixed for 12 minutes in 5% methanol-free formaldehyde. Ovaries underwent four 10-minute washes in 0.5 mL PBT (1X PBS, 0.5% Triton X-100, 0.3% BSA) while nutating. Primary antibodies in PBT were applied and incubated overnight at 4°C with nutation. Samples were then washed three times for 5-8 minutes each in 1 mL PBT. Secondary antibodies were added in PBT with 4% donkey serum and incubated at room temperature for 2-3 hours. Samples underwent three 10-minute washes in 1 mL of 1X PBST (0.2% Tween 20 in 1x PBS) and were incubated in Vectashield with DAPI (Vector Laboratories) for a minimum of 1 hour before mounting.

### Antibodies

Primary antibodies utilized were: mouse anti-1B1 (1:20; DSHB), Rabbit anti-Vasa (1:1,000; Rangan Laboratory), Chicken anti-Vasa (1:1,000; Rangan Laboratory), Rabbit anti-GFP (1:2,000; abcam, ab6556), Rabbit anti-H3K9me3 (1:500; Active Motif, AB_2532132), Mouse anti-H3K27me3 (1:500; abcam, ab6002), Rabbit anti-Egl (1:1,000; Lehmann Laboratory), Mouse anti-NPC (1:2000; BioLegend, AB_2565026). Secondary antibodies (Alexa 488 from Molecular Probes, Cy3 and Cy5 from Jackson Labs) were used at a 1:500 dilution.

### Fluorescence Microscopy

Ovaries were mounted on slides and imaged using Zeiss LSM-710 and LSM-880 confocal microscopes under 20X, 40X and 63X oil objectives with pinhole set to 1 airy unit. Image processing was conducted using Fiji, with gain adjustment and cropping performed in Photoshop.

### RNA Extraction and TURBO DNase Treatment

Ovaries were dissected into PBS, transferred to RNase-free microcentrifuge tubes, and flash-frozen in 100μl Trizol at -80°C. Samples were lysed using a plastic disposable pestle, with Trizol added to 1mL total volume. After 5 minutes at room temperature, samples were centrifuged (20 min, >13,000g, 4°C). The supernatant was mixed with 500μl chloroform, incubated for 5 minutes at room temperature, and centrifuged (10 min, max speed, 4°C). The aqueous phase underwent ethanol precipitation with sodium acetate (10% of transferred volume) and 2-2.5 volumes of 100% ethanol at -20°C overnight. RNA was pelleted by centrifugation (15 min, max speed, 4°C), washed with 75% ethanol, dried, and resuspended in 20-50μl RNase-free water. Concentration was measured via nanodrop (Blatt et al., 2021).

### RNA-seq Library Preparation and Analysis

Libraries were prepared using the Revvity NEXTFLEX Rapid Directional RNA-Seq Kit 2.0 (NOVA-5198-02). RNA was treated with Turbo DNase (TURBO DNAfree Kit, Life Technologies, AM1907) at 37°C for 30 min. DNase was inactivated, and RNA was purified by centrifugation. Poly-A selection was performed on normalized RNA quantities. mRNA libraries were constructed, with quantity assessed via Qubit and quality via Bioanalyzer or Fragment Analyzer. Sequencing generated single-end 100 base pair reads on an Illumina NextSeq500.

Bioinformatic analysis employed FastQC (v0.11.8), Trim Galore! (v0.6.6), STAR aligner (v2.7.5b), and Salmon (v1.2.1). Differential expression analysis utilized the DESeq2 R package (v1.30.1). Genes with <5 total reads across all samples were filtered out. Differential expression criteria: adjusted p-value <0.05 and |log_2_ fold change| ≥2. Plots were generated using ggplot2 in R (v.4.3.1) (Kotb et al., 2024).

### DNA FISH with Immunofluorescence

Ovaries were fixed, washed, and incubated with primary antibodies overnight. After secondary antibody incubation, samples underwent a series of washes with increasing formamide concentrations. Pre-denaturation was performed at varying temperatures, followed by probe hybridization overnight at 37°C. Samples were washed, mounted in Vectashield, and imaged (Sarkar et al., 2023).

### CUT & RUN Assay

Ovaries were permeabilized, incubated with antibodies overnight, and treated with pAG-MNase. DNA cleavage was induced with CaCl2, followed by RNase treatment and Proteinase K digestion. DNA was purified using magnetic beads and quantified via Qubit assay and Fragment analyzer.

DNA-seq Library Preparation and Analysis Libraries were prepared using the NEBNext Ultra II DNA Library Prep Kit. Bioinformatic analysis employed FastQC, Trim Galore!, Bowtie2, samtools, MACS2, deepTools, ChipSeeker, HOMER, and ChromHMM. Various genomic analyses and visualizations were performed using these tools (Kotb et al., 2024).

### Polysome seq

Polysome profiling of ovaries was carried out with modifications based on established protocols (44). In brief, we dissected ovaries from approximately 100 YWT flies (UAS-Dcr2; nosGAL4) or around 275 pairs of ovaries from *zfrp8* experimental flies using PBS. The dissected ovaries were immediately flash-frozen in liquid nitrogen. The frozen tissue was then homogenized in lysis buffer using a motorized pestle, with 20% of the lysate reserved for mRNA extraction and library preparation as previously described. The remaining lysate was loaded onto sucrose gradients (10–45%) supplemented with cycloheximide, using Beckman Coulter 9 of 16 × 3.5 PA tubes (#331372), and centrifuged at 35,000g in an SW41 rotor for 3 hours at 4°C. After centrifugation, the gradients were fractionated using a density gradient fractionation system. RNA was extracted from the fractions with acid phenol–chloroform, precipitated overnight, and the resulting pellet was resuspended in 20 μl of water, treated with TURBO DNase, and used for library preparation as outlined above (McCarthy et al., 2022).

## Data Availability

Raw and unprocessed RNA-seq and CUT & RUN data are available in the Gene Expression Omnibus (GEO) databank under accession number GEOXXXX.

## Acknowledgements

We thank members of the Rangan laboratory and Dr. Vernon Monteiro for their comments on the manuscript. We thank the Bloomington *Drosophila* Stock Center, Vienna *Drosophila* Resource Center, Berkeley *Drosophila* Genome Project Gene Disruption Project, and FlyBase for reagents and resources. P.R. is funded by the NIH/NIGMS (R35GM152236). This work was supported in part by the Bioinformatics for Next-Generation Sequencing (BiNGS) shared resource facility within the Tisch Cancer Institute at the Icahn School of Medicine at Mount Sinai, which National Institutes of Health grant P30CA196521 partially supports; the Black Family Stem Cell Institute; and the Department of Cell, Developmental, and Regenerative Biology at the Icahn School of Medicine at Mount Sinai. This work was also supported in part through the computational resources and staff expertise provided by Scientific Computing and Data at the Icahn School of Medicine at Mount Sinai and supported by the Clinical and Translational Science Awards (CTSA) grant UL1TR004419 from the National Center for Advancing Translational Sciences. The research reported here was supported by the Office of Research Infrastructure of the National Institutes of Health under award number S10OD026880.

---

**Supplementary Figure 1.**
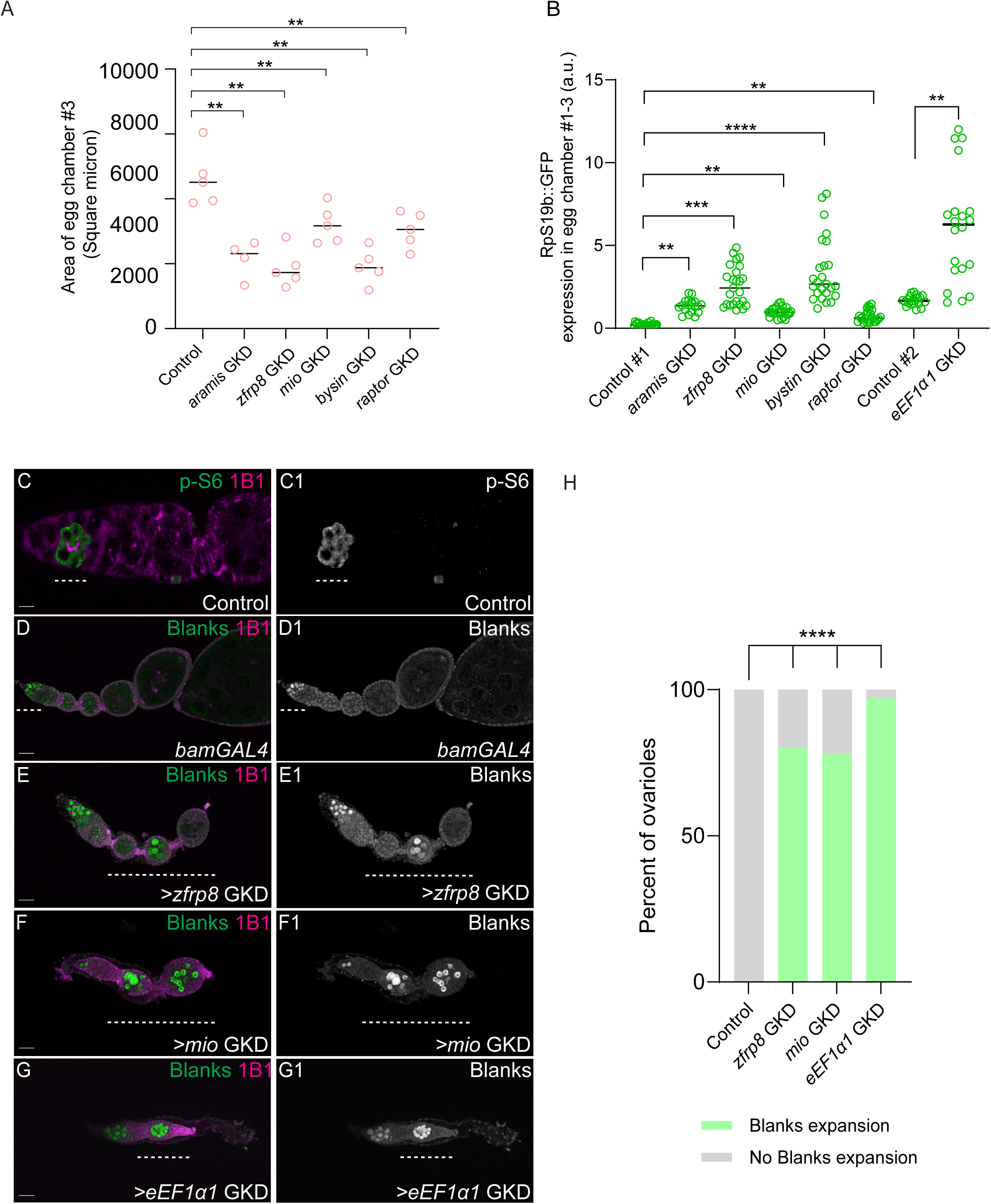
Ribosome level regulators are required in the cyst stages for silencing early oogenesis gene *blanks* during oogenesis. (A) Quantification of the size of the egg chambers of control and *zfrp8* GKD, *bystin* GKD, *mio* GKD, *raptor* GKD and *eEF1α1* GKD ovaries. Graph shows that *zfrp8* GKD, *bystin* GKD, *mio* GKD, and *raptor* GKD’s egg chambers do not grow compared to control ovaries. *eEF1α1* GKD did not lead to the formation of a third egg chamber therefore we were unable to quantitate. Statistics: Two-tailed t-test; n = 5 ovarioles per genotype; ns, P > 0.05; ∗p < 0.05; ∗∗p < 0.01; ∗∗∗p < 0.001; ∗∗∗∗p < 0.0001. (B) Arbitrary unit (A.U.) quantification of early oogenesis proteins RpS19b::GFP expression in *zfrp8* GKD, *bystin* GKD, *mio*GKD, *raptor* GKD and *eEF1α1* GKD compared to control ovaries. RpS19b::GFP persists in egg chambers of *zfrp8*GKD, *bystin* GKD, *mio* GKD, *raptor* GKD and *eEF1α1* GKD compared to the egg chambers of control. n = 5 ovarioles per genotype. Statistics: Two-tailed t-test; n = 50 ovarioles per genotype; ns, P > 0.05; ∗p < 0.05; ∗∗p < 0.01; ∗∗∗p < 0.001; ∗∗∗∗p < 0.0001. (C-C1) Wild type control germarium (A) and control grayscale (B) stained for p-S6 (green, right in grayscale) and 1B1 (magenta). p-S6 marking TOR activity is expressed in the cyst stages. (D-D1) Ovariole of control *bamGAL4;nosGAL4* (B) and grayscale (B1) stained for blanks (green, right in grayscale) and 1B1 (magenta). In the control, blanks is expressed in the undifferentiated cells (yellow arrows) and is attenuated in the egg chambers. (E-E1) Ovariole of *bamGAL4*;*nosGAL4> zfrp8* GKD (C) and grayscale (C1) stained for blanks (green, right in grayscale) and 1B1 (magenta). *zfrp8* GKD in the cyst stages using *bamGAL4*;*nosGAL4* resulted in egg chambers that ectopically expressed blanks (white dashed line), did not grow, and died during oogenesis. (F-F1) Ovariole of *bamGAL4*;*nosGAL4> mio* GKD (D) and grayscale (D1) stained for blanks (green, right in grayscale) and 1B1 (magenta). *mio* GKD in the cyst stages using *bamGAL4*;*nosGAL4* resulted in egg chambers that ectopically expressed blanks (white dashed line), did not grow, and died during oogenesis. (G-G1) Ovariole of *bamGAL4*;*nosGAL4> raptor* GKD (E) and grayscale (E1) stained for blanks (green, right in grayscale) and 1B1 (magenta). *raptor* GKD in the cyst stages using *bamGAL4*;*nosGAL4* resulted in egg chambers that ectopically expressed blanks (white dashed line), did not grow, and died during oogenesis. (H) Quantitation of percent ovarioles with Blanks expansion in control, *zfrp8* GKD, *bystin* GKD, *mio* GKD, *raptor* GKD and *eEF1α1* GKD ovaries. Statistics: Two-tailed t-test; n = 50 ovarioles per genotype; ns, p > 0.05; ∗p < 0.05; ∗∗p < 0.01; ∗∗∗p < 0.001; ∗∗∗∗p < 0.0001. Scale bars: (C-C1) 7.5 μm; (D-G1) 15 μm.

**Supplementary Figure 2.**
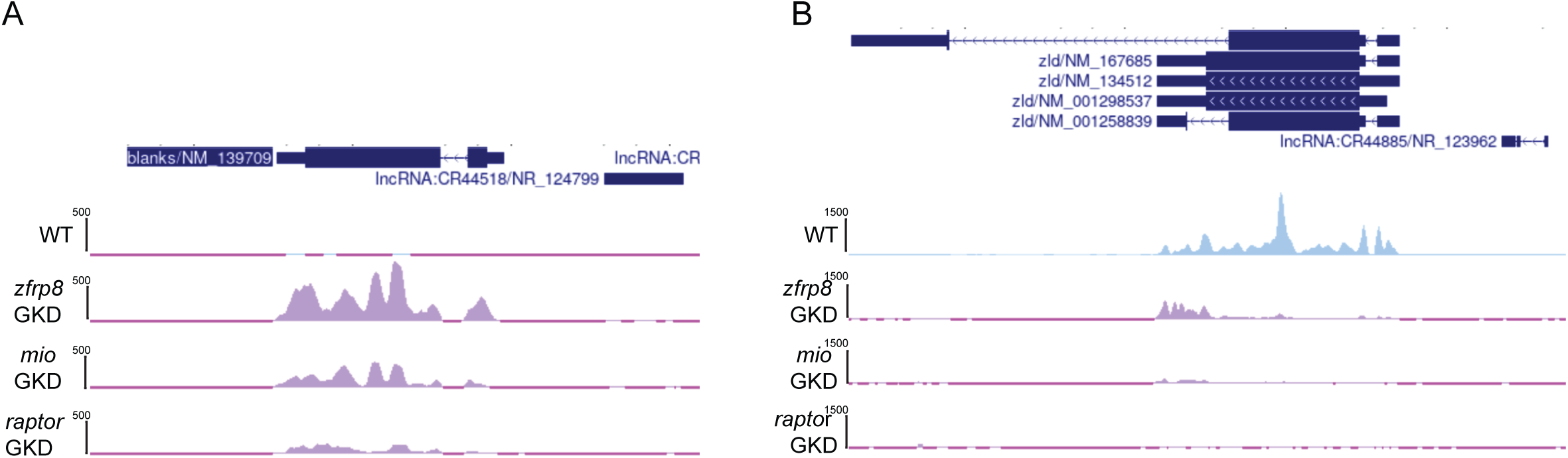
Ribosome level regulators are required for silencing early oogenesis genes during oogenesis. (A) RNA-seq tracks showing that *blanks* is upregulated upon >*zfrp8* GKD, >*mio* GKD and >*raptor* GKD (purple) compared to control *nosGAL4* (blue). (B) RPKM-normalized RNA-seq tracks showing that *zelda* is downregulated upon >*zfrp8* GKD, >*mio* GKD and >*raptor* GKD (purple) compared to control (*nosGAL4)* (blue).

**Supplementary Figure 3.**
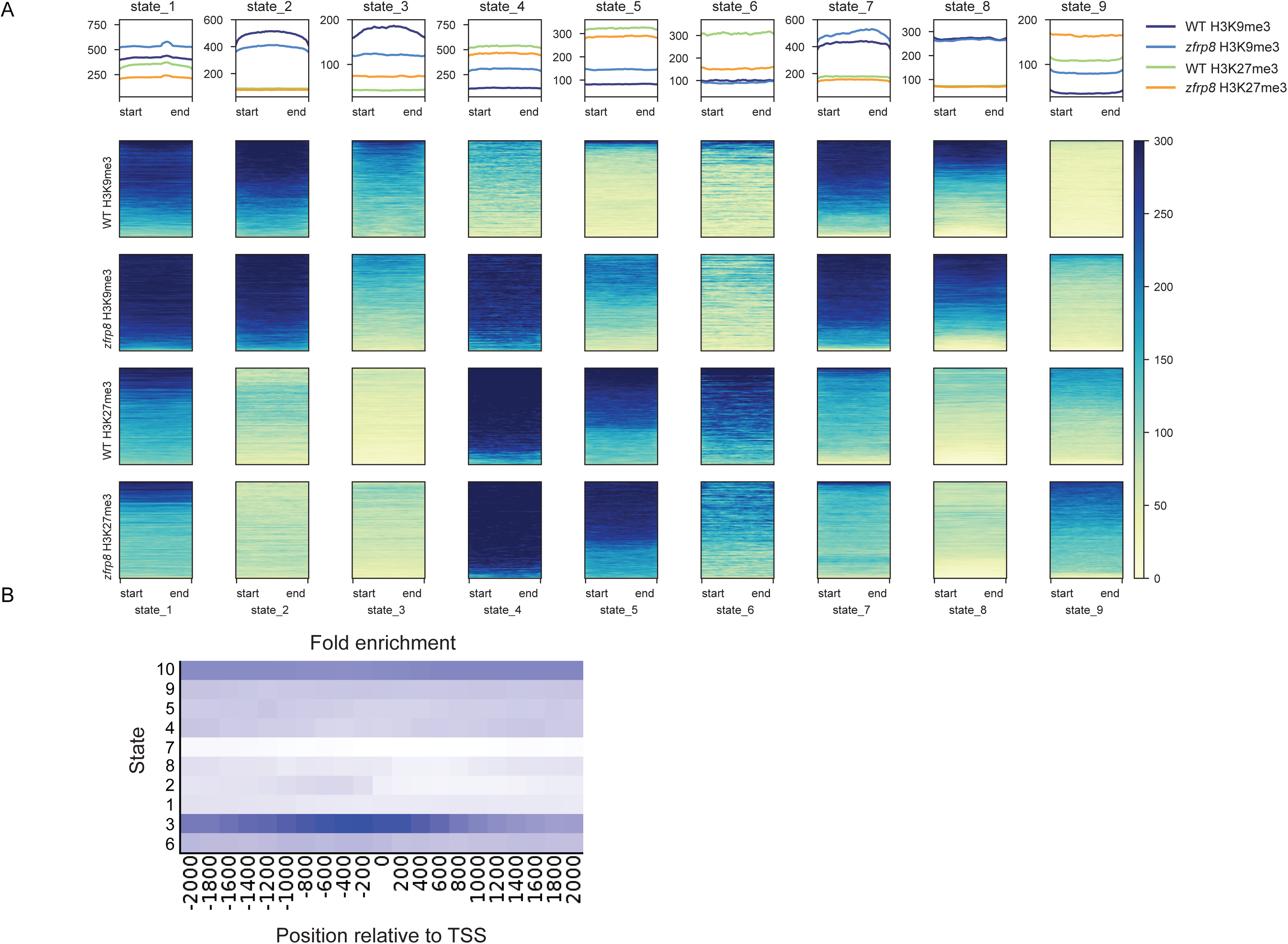
Ribosome level regulators are required maintaining proper chromatin state. (A) A 10-state chromatin model depicting H3K9me3 and H3K27me3 distribution across the genome. This model illustrates H3K9me3 and H3K27me3 dynamics across 10 chromatin states, highlighting their distribution in different genomic regions. In *zfrp8* GKD, H3K9me3 is reduced on promoters in state 3. These findings reveal that ribosome biogenesis influences genome-wide epigenetic landscapes, coordinating transcriptional regulation during oogenesis. (B) In state 3, H3K9me3 is enriched around promoters.

**Supplementary Figure 5.**
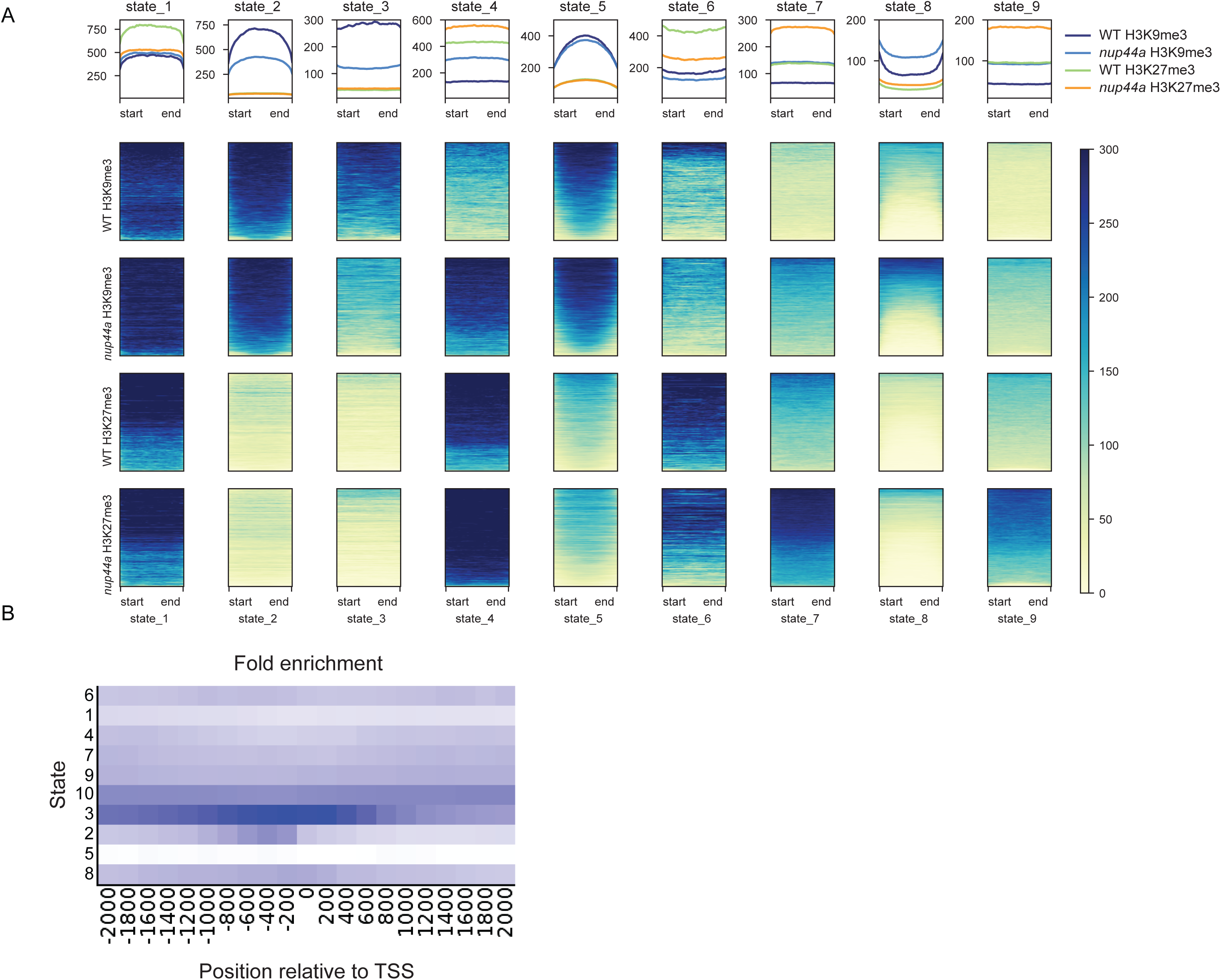
Ribosome level regulators are required for maintaining proper chromatin state. (A) A 10-state chromatin model depicting H3K9me3 and H3K27me3 distribution across the genome. This model illustrates H3K9me3 and H3K27me3 dynamics across 10 chromatin states, highlighting their distribution in different genomic regions. In *Nup44A* GKD, H3K9me3 is reduced on promoters in State 3 and H3K27me3 is redistributed across different states, indicating widespread chromatin reorganization. These findings reveal that Nup44A influences genome-wide epigenetic landscape. (B) In state 3, H3K9me3 is enriched around promoters, ensuring transcriptional repression of genes including *rps19b*.

## Tables and their legends

**Supplementary Table 1: RNA-seq analysis of differentially expressed genes in *zfrp8*, *mio*, and *raptor* GKD ovaries.** This table lists genes that are significantly upregulated or downregulated in GKD ovaries compared to controls (|log₂FC|) ≥ 2, FDR < 0.05). Upregulated genes include early oogenesis transcripts, whereas downregulated genes include maternal RNAs such as *wispy* and *zelda*. Shared and unique targets across the three GKD conditions are indicated. These findings suggest that ribosome biogenesis and TORC1 activity regulate gene expression during oocyte specification by modulating RNA stability and/or transcription.

**Supplementary Table 2: Genes co-regulated by ribosome biogenesis and chromatin modifiers** This table lists genes that are upregulated in *zfrp8*, *mio*,and *raptor* GKD ovaries compared to controls. Genes were identified using RNA-seq analysis with a log_2_FC and an FDR < 0.05. To determine whether ribosome biogenesis regulators share targets with chromatin modifiers, we compared these upregulated genes with previously published datasets of genes misregulated in *SETDB1*, *Nup154*, and *e(z)* GKD ovaries. Genes that overlap between ribosome regulators and chromatin regulators are highlighted, representing a shared set of early oogenesis genes that require both translation and chromatin remodeling for proper repression during oocyte specification.

**Supplementary Table 3. List of translationally upregulated and downregulated genes in *zfrp8* GKD ovaries identified by polysome-seq.** The table includes gene names, log_2_FC and FDR values compared to control ovaries. Genes that are translationally upregulated include those involved in siRNA-mediated heterochromatin formation and chromatin organization (*Su(Hw)*, *M1BP*), suggesting a potential feedback response to heterochromatin loss. Translationally downregulated genes include ribosomal components, *Nup44A,* a nuclear pore complex protein, and *lola-like*/*batman*, a BTB/POZ domain protein involved in chromatin regulation.

**Supplementary Table 4: Differentially expressed genes in *Nup44A* GKD ovaries and their comparison to ribosome and chromatin regulators.** This table presents RNA-seq analysis of genes differentially expressed in *Nup44A* GKD ovaries compared to controls. Differential expression was determined using an |log₂FC| ≥ 2 and an FDR < 0.05. The table includes upregulated and downregulated genes in *Nup44A* GKD and their overlap with targets misregulated in *zfrp8*, *mio*, and *raptor* GKD (ribosome biogenesis/TORC1 regulators) as well as *SETDB1*, *Nup154*, and *e(z)* GKD (chromatin regulators).

