## Supplementary figures and images for "TORC1-driven translation of *Nucleoporin44A* promotes chromatin remodeling and germ cell-to-maternal transition in Drosophila"

### Supplementary Figures

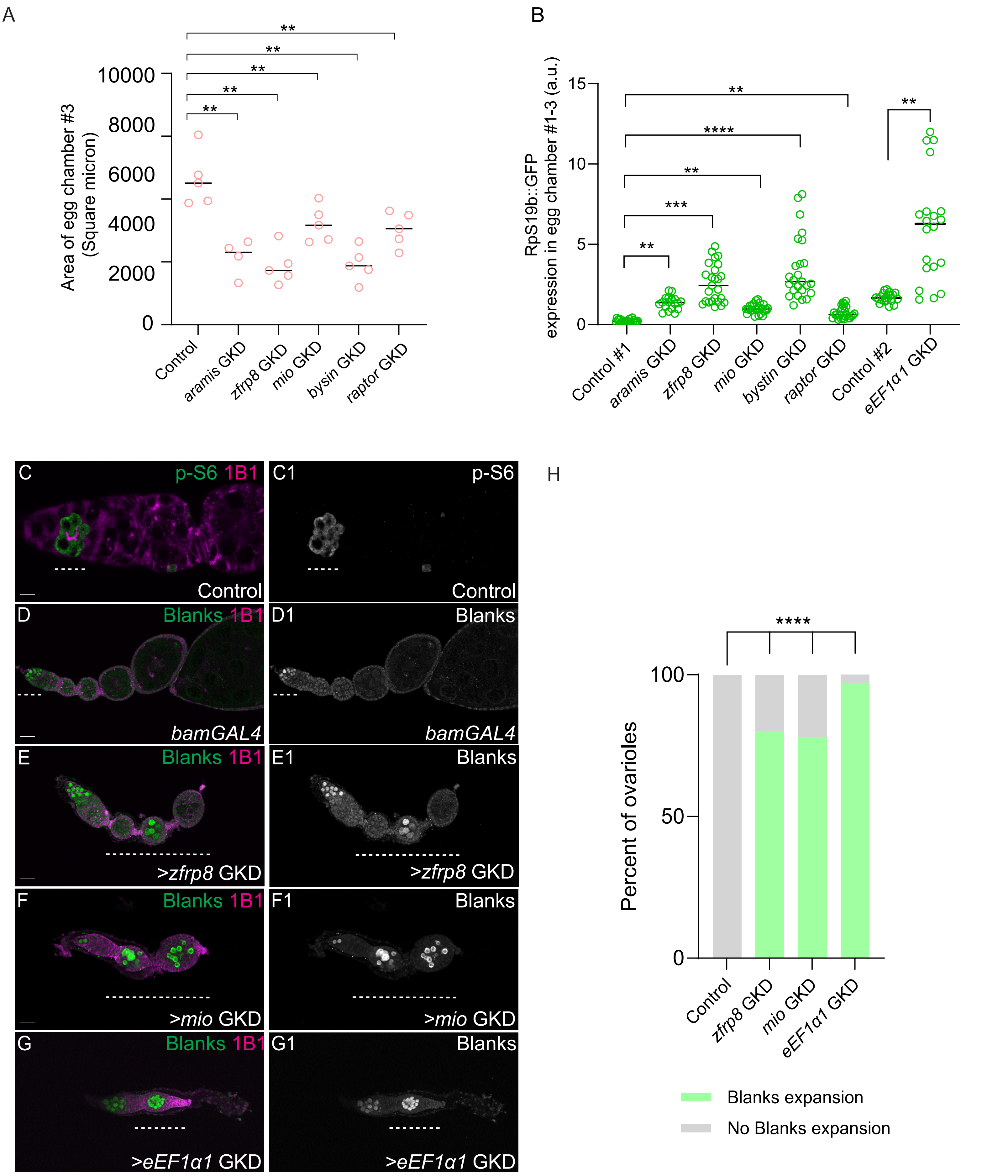

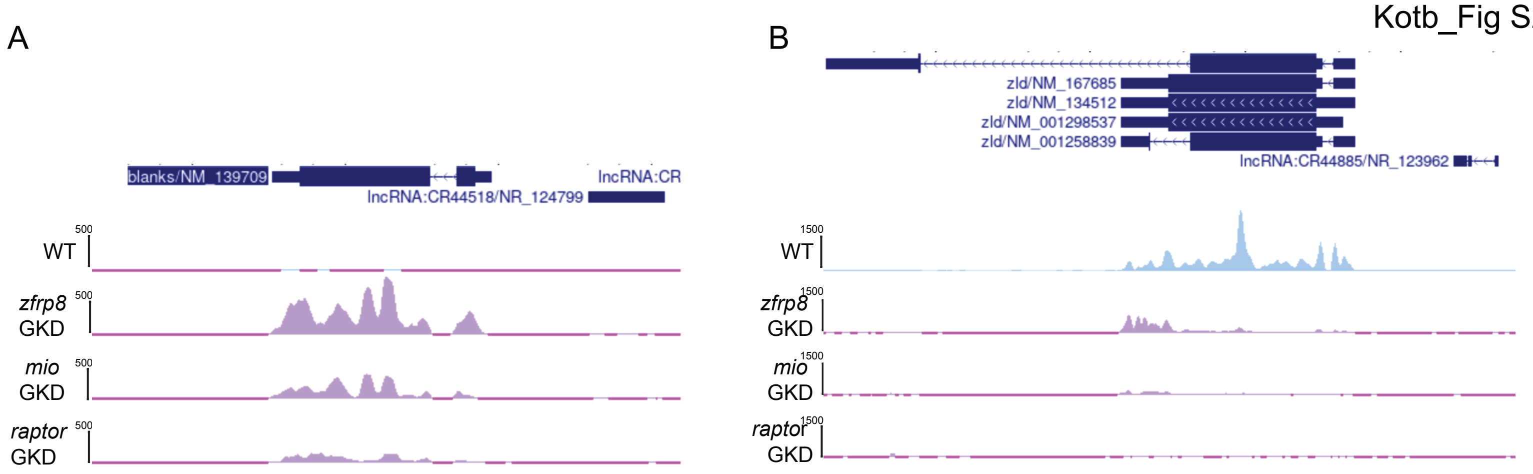

Kotb_Fig S3

A

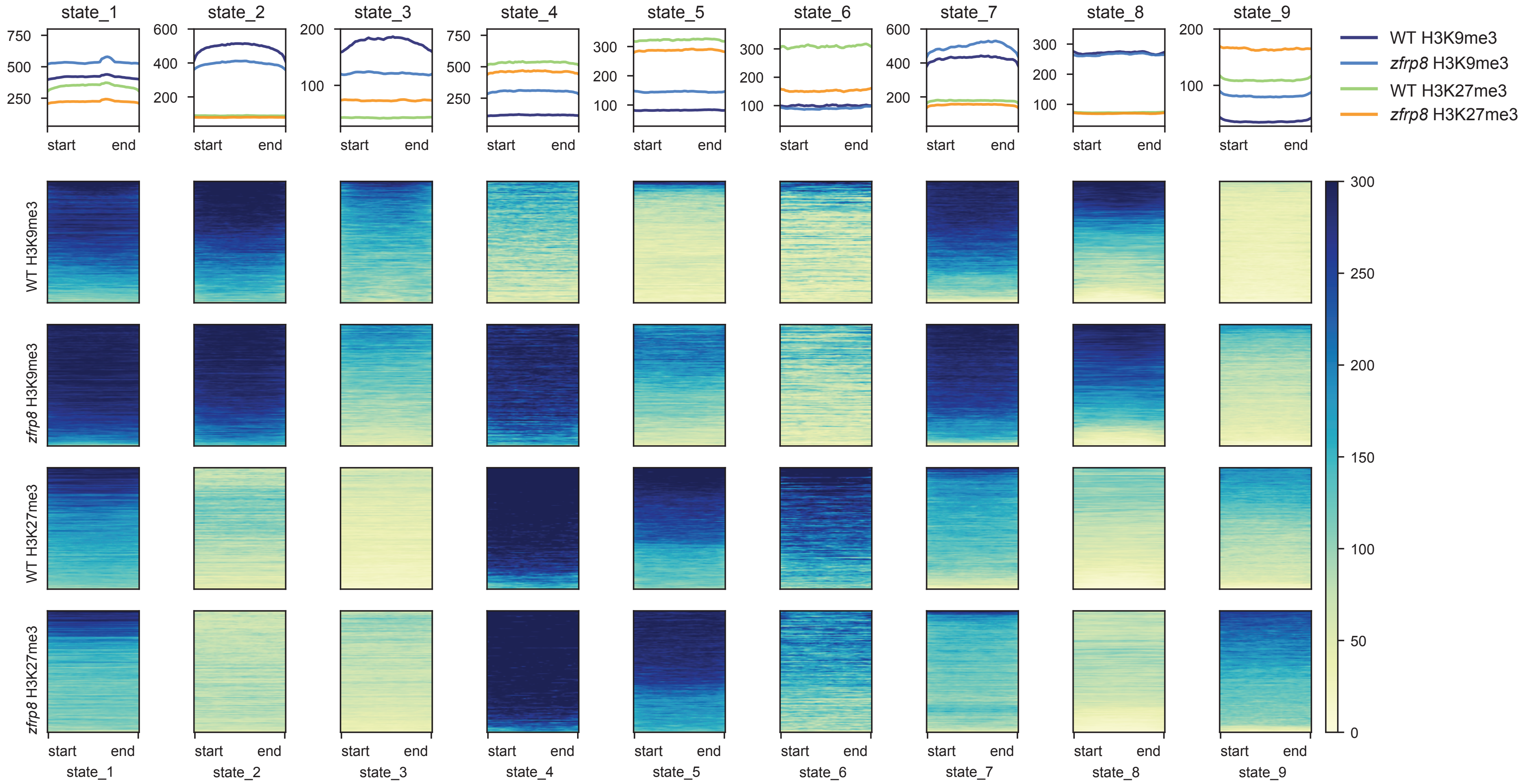


B

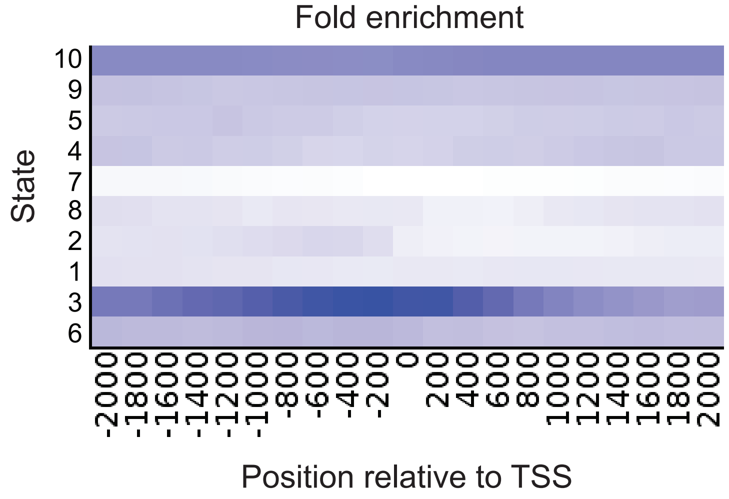

Kotb_Fig S5

A

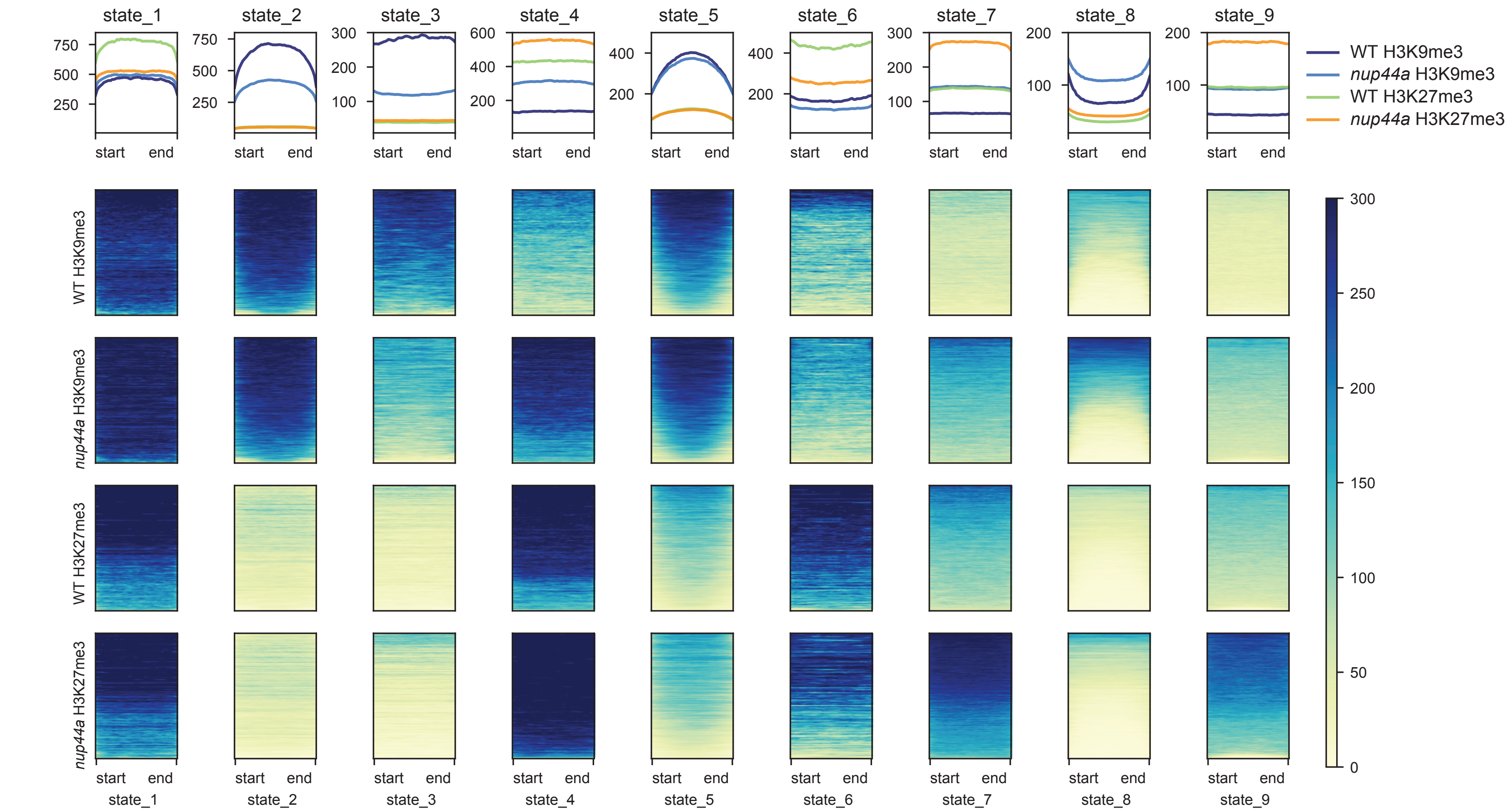


B

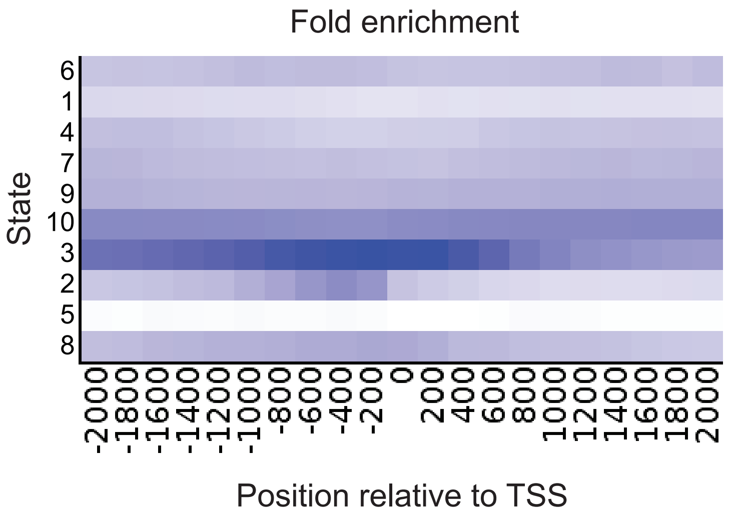
